# Urban Vietnam at the microbiome transition: Co-existing enterotypes reflect the global industrialization patterns

**DOI:** 10.64898/2026.10.08.757770

**Authors:** Thanh Thien Le, Joshua Tran, Linh Hong Thi Mai, Dao Bich Thi Tran, Dat Quoc Nguyen, Quynh Mai Nguyen, Giang Minh Vu, Minh Ngoc Truong, Manh Khanh Dinh, Hao The Chung, Thanh Hong Nguyen, Chi Du Thi Tran, Huan Duc Pham, Huong Thu Thi Pham, Muoi Xuan Nguyen, Thuy Xuan Nguyen, Hang Dao Viet, May Ngoc Le, Tuan Thanh Nguyen, Jennifer Le, Jack Gilbert, Xuan-Hung Nguyen

**Affiliations:** Vinmec-VinUni Institute of Immunology, College of Health Sciences, VinUniversity, Hanoi, Vietnam; Innovations in Health Sciences, VinUniversity, Hanoi, Vietnam; Vinmec International Hospital, Hanoi, Vietnam; Department of Pediatrics, University of California San Diego, La Jolla, CA, US; Center for Marine Biotechnology and Biomedicine, Scripps Institution of Oceanography, La Jolla, CA, USA; Discipline of Surgery, School of Medicine, College of Health, Adelaide University, Adelaide, SA, Australia; Department of Surgery - Otolaryngology Head and Neck Surgery and the Basil Hetzel Institute for Translational Health Research, The Queen Elizabeth Hospital, Central Adelaide Local Health Network, Adelaide, SA, Australia; Oxford University Clinical Research Unit, Ho Chi Minh City, Vietnam; Saw Swee Hock School of Public Health, National University of Singapore, Singapore, Singapore; Center for Gastroenterology - Hepatobiliary – Urology, Vinmec Times City, Vinmec International Hospital, Hanoi, Vietnam; College of Health Sciences, VinUniversity, Hanoi, Vietnam; Internal Medicine Faculty, Hanoi Medical University, Hanoi, Vietnam; Endoscopy Center, Hanoi Medical University Hospital, Hanoi, Vietnam; Department of Oncology, Vinmec Times City, Vinmec International Hospital, Hanoi, Vietnam; Department of Pharmacy Practice and Sciences, University of California San Diego Skaggs School of Pharmacy and Pharmaceutical Sciences, La Jolla, CA, USA; Center for International Pharmacy Education and Research, University of California San Diego, La Jolla, CA, USA

## Abstract

Vietnam’s rapid socioeconomic and dietary transition offers a rare window into how industrialization is reflected in the gut microbiome, yet Vietnamese populations remain underrepresented in global reference datasets. We investigated whether healthy Vietnamese adults in Hanoi harbor microbial community configurations associated with different stages of industrialization. Metagenomic data from stool samples of 183 clinically screened participants in Hanoi were analyzed alongside 629 publicly available metagenomic datasets from populations in low to middle income (LMI) and high income (HI) countries. Species-level taxonomic composition and gene-content-based metabolic pathways were characterized using community clustering and multivariable association analyses. Rather than forming a single population profile, the Vietnamese cohort separated into two enterotypes that mirrored the global industrialization gradient. Enterotype 1 more closely resembled populations from LMICs and showed greater microbial richness and diversity, whereas Enterotype 2 aligned with taxonomic and functional profiles observed in HICs. Both taxonomic and functional potential shows similar trajectories. Diet and lifestyle patterns describe some of the variance between enterotypes, supporting an association between contemporary exposures and microbiome structure without establishing causation. Antibiotic resistance profiles also varied between enterotypes, suggesting that the transition may encompass functions relevant to antimicrobial exposure. Together, these findings position urban Vietnam at a microbiome transition, where community states associated with different levels of industrialization coexist within a single population. This Vietnamese reference cohort expands representation in global metagenomic research and provides a baseline for tracking how rapid socioeconomic change may reshape microbial ecology, metabolic capacity, and antimicrobial resistance.

## Introduction

The human gut microbiome plays a fundamental role in host metabolism, immune development, nutrient utilization, and resistance to disease^1,2^. Increasing evidence demonstrates that microbial communities are strongly shaped by geography, diet, lifestyle, urbanization, and environmental exposures, making the concept of a universal “healthy microbiome” increasingly untenable^3,4^. Consequently, population-specific reference microbiomes have become an essential foundation for precision medicine^1,4^, enabling accurate interpretation of disease-associated dysbiosis^2^ and the development of microbiome-based diagnostics and therapeutics. Despite substantial advances through international microbiome initiatives, current reference datasets remain heavily biased toward populations from China, Europe, and North America^5,6^. Large-scale metagenomic studies in China, encompassing thousands of fecal samples, have substantially expanded microbial reference genome catalogs and characterized population-associated genomic variation^7–9^. In contrast, Southeast Asian populations remain relatively underrepresented despite experiencing some of the most rapid socioeconomic and dietary transitions^10^. This imbalance limits our understanding of global microbiome diversity^6,11^ and raises important questions about how microbial ecosystems respond to changing lifestyles^11,12^.

Vietnam presents a particularly informative natural model for addressing these questions. Historically, Vietnam has maintained one of the lowest adult obesity prevalences worldwide, with an estimated prevalence of only approximately 2.1% in 2022^13^. However, rapid economic development, urbanization, and westernization over the past two decades have substantially altered dietary patterns, physical activity, and environmental exposures^14,15^, resulting in a rapidly increasing prevalence of overweight and obese phenotypes^16^. Rather than representing a stable population, modern Vietnam encompasses individuals living across a broad spectrum of traditional and westernized lifestyles^14^, offering a unique opportunity to investigate how gut microbial communities adapt during epidemiological and nutritional transition.

Beyond these metabolic changes, Vietnam also represents an important setting for studying microbiome ecology because of its exceptionally high burden of antimicrobial exposure^17,18^. Extensive antibiotic use in both human medicine and agriculture has contributed to one of the largest known reservoirs of antimicrobial resistance genes in healthy individuals^17^. Consequently, the Vietnamese gut microbiome reflects not only changing dietary and lifestyle patterns but also long-term environmental selection pressures that may shape both microbial community structure and functional potential^4,15^. Understanding the baseline microbiome in this context is therefore important not only for metabolic health but also for understanding the distribution of the gut resistome within healthy populations^17^.

Together, these characteristics make Vietnam an interesting population in which to establish a healthy reference microbiome. Unlike relatively stable populations that largely represent either traditional or Westernized lifestyles, Vietnam is undergoing rapid transition and therefore provides a unique opportunity to investigate how healthy gut microbial communities are organized during this process. We hypothesized that the healthy Vietnamese gut microbiome would not comprise a single homogeneous community but instead contain multiple ecological configurations reflecting different stages along the transition between traditional and urbanized microbial compositions.

In this project, we generated one of the largest metagenomic datasets of healthy Vietnamese adults and compared it with publicly available healthy cohorts representing diverse geographic regions and dietary backgrounds. Using taxonomic profiling, ecological network analysis, diversity analyses, and unsupervised clustering, we examined both within-population heterogeneity and between-population relationships. We identified two microbiome configurations within the healthy Vietnamese population that consistently aligned with microbial community structures characteristic of low-to-middle income countries (LMICs) and high-income countries (HICs), respectively. These findings establish a population-specific reference microbiome for Vietnam while suggesting that populations undergoing rapid lifestyle transition may simultaneously harbour multiple stable ecological states that bridge rural-traditional and urbanized-modern gut microbiomes.

## Methods

### Participant Recruitment

Stool samples were obtained from two sources: archived samples from the VinCAPR program^19^ and samples from participants newly recruited for the present study. VinCAPR offered population-based free screening to adults aged ≥40 years across six Hanoi districts, four of which were classified as suburban or rural. Newly recruited participants came primarily from the Hanoi metropolitan area. The same inclusion and exclusion criteria for the present analysis were applied to participants from both sources.

Eligible participants were 20–80 years of age, had a body mass index (BMI) between 18 and 30 kg/m², normal vital signs, and routine blood tests within the reference ranges. Participants were also required not to follow a specialized or restrictive diet, to consume fewer than seven servings of alcohol per week, and to have no colorectal polyps detected during colonoscopy. Exclusion criteria included chronic infectious or immune-mediated diseases, including human immunodeficiency virus infection, hepatitis, syphilis, and autoimmune disease. Individuals were also excluded if they had used antibiotics, immunosuppressive agents, or proton pump inhibitors, or had received any vaccination within the preceding three months. Additional exclusion criteria included a history of cancer, diabetes mellitus, cardiovascular disease, or stroke; gastrointestinal conditions such as inflammatory bowel disease, irritable bowel syndrome, gastroenteritis, chronic diarrhea, or chronic constipation; and previous gastrointestinal surgery.

For newly recruited participants, an initial telephone screening was conducted to assess general health status and preliminary eligibility. Individuals who passed the telephone screening underwent a comprehensive health examination, including physical assessment, routine blood tests, and colonoscopy, at Vinmec Times City International Hospital. Final eligibility was determined according to predefined inclusion and exclusion criteria. All participants provided written informed consent before enrollment. The study protocol was approved by the Institutional Review Board of Vinmec International Hospital (approval number: 83/2023/CN/HĐĐĐ VMEC).

### Sample collection and processing

Participants received kits for stool self-collection. VinCAPR samples were collected on stool collection cards using Fe-Col collection paper (FC2010, Alpha Laboratories) or in OC-Sensor tubes (Eiken Chemical). Samples were kept at 2–8°C until pick-up and transported to −80°C for long-term storage. Samples from newly recruited participants were collected in DNA/RNA Shield preservative (Zymo Research), stored at 4°C after collection, and transferred to −80°C at the laboratory.

The same downstream DNA extraction, library preparation, and sequencing protocols were used for samples from both recruitment sources. Microbial DNA was extracted from stool samples using the QIAamp PowerFecal DNA Kit (QIAGEN), including bead-assisted mechanical disruption, according to the manufacturer’s instructions. Libraries sequenced on an Illumina NextSeq 2000 platform, with a target sequencing depth of approximately 20 million PE150 read pairs per sample. Samples were sequenced with environmental and Zymo mock communities controls.

### Public dataset

Publicly available metagenomic datasets were initially identified through the Microbiome Atlas and published literature. Datasets were selected to provide broad geographic representation across continents and to include populations from both HI and LMI countries. Eligible datasets were required to provide participant-level metadata on age, sex, and BMI. Samples from adults older than 18 years with a BMI between 18 and 30 kg/m² were included, with the exception of Tanzania (no BMI). Where possible, samples were selected to achieve a balanced distribution of age and sex across countries. Only one sample per participant was used. Details of the original studies, BioProject accession numbers, and sequence accession numbers are provided in Supplementary Table 1 and Supplementary File 1.

## Data Analysis

### Preprocessing, taxonomic and functional profiling

All metagenomes underwent the same preprocessing, taxonomic, and functional profiling. In short, paired-end reads were adapter-trimmed and quality-filtered with fastp v0.22.0^20^, host and control reads were removed with minimap2 v2.22^21^. SAMtools v1.15.1^22^ retained only pairs with both mates unmapped. Species-level relative abundances were estimated with MetaPhlAn v4.1.1^23^, using the mpa_vJun23_CHOCOPhlAnSGB_202403 marker database and Bowtie2 v2.5.4 for read mapping^24^. Encoded functional potential was profiled with HUMAnN v3.9^25^. Unmapped reads underwent translation alignment against UniRef90 (2019_01b) using DIAMOND v2.1.11^26^. Co-abundance networks were inferred using SPIEC-EASI and StARS^27^, retaining species with prevalence ≥20% and mean relative abundance >0.1%. Antibiotic resistance genes (ARGs) were profiled using Resistance Gene Identifier (RGI)^28^ and matched to WHO priority antibiotic-resistant pathogens^29^.

### Differential signature identification

Differential species abundance and prevalence were assessed with MaAsLin3 v1.4.0^30^, retaining features with prevalence ≥10% and mean relative abundance ≥0.01%. Models included geographic group, age, gender and read depth as fixed effects, with Vietnam and female as reference categories and no random effects. Relative abundances underwent total-sum scaling and log transformation; age and read depth were z-score standardized. P values were Benjamini–Hochberg corrected, with significance defined as joint q ≤0.1. To identify taxa consistently distinguishing Vietnam, species-specific coefficients were aggregated across the five group-versus-Vietnam contrasts using q <0.05.

### Enterotype clustering

Vietnamese samples were clustered using Dirichlet–multinomial mixture (DMM) modeling^31^. Genus-level counts were filtered to retain genera present in ≥10% of samples and rarefied to 10,000 reads per sample. Models with K = 1–6 components were evaluated using the Laplace approximation, Akaike information criterion and Bayesian information criterion.

### Industrialization score

Industrialization-associated scores were derived from species and MetaCyc pathway profiles in public cohorts using elastic-net logistic regression^32^ and coefficient-weighted MaAsLin3 models^30^. Elastic-net performance was evaluated by nested leave-one-study-out cross-validation. Models and preprocessing parameters were applied unchanged to Vietnamese samples. Scores were rescaled without truncation to reference medians of 0 (non-industrialized) and 1 (industrialized). Enterotypes were compared using bootstrap estimates and Wilcoxon/Welch’s tests. Covariate-adjusted global and Vietnamese feature effects were compared using Spearman correlation; features significant in both models (BH-adjusted q <0.05) were classified as concordant or discordant by effect direction.

### Lifestyle associations and enterotype classification

Twelve dietary and lifestyle variables (diet score, seven dietary components, smoking, walking, regular exercise and weekly alcohol use) were tested separately for association with Vietnamese enterotypes using logistic regression adjusted for age, sex and BMI. P values were Benjamini–Hochberg corrected across the 12 tests. Variables selected from these analyses were used in an L2-penalized logistic classifier, tuned and evaluated by repeated stratified five-fold cross-validation (20 repeats). Models incorporating all 12 variables, including L1- and L2-penalized logistic regression, random forest and XGBoost, were evaluated using the same procedure. All classifiers used class-balanced weights; performance was summarized as mean ± SD ROC-AUC across repeats.

## Results

### Two community configurations differ in richness and diversity

Participant selection is summarized in Supplementary Figure S1A. After quality control, the analyzed cohort comprised 183 healthy adults in Hanoi, including 108 women and 75 men, with 86 newly recruited participants and 97 VinCAPR participants (Fig. 1A–C; Supplementary Fig. S1A). VinCAPR participants were older than newly recruited participants (*P* < 0.001), whereas BMI and sex distributions did not differ significantly (*P* = 0.347 and 0.200, respectively; Supplementary Fig. S1B–D). Sequencing quality was high across recruitment sources and batches (Supplementary Fig. S1E,F). Samples from the two sources occupied overlapping regions of the principal coordinates analysis (PCoA) plot, and recruitment source accounted for 3.4% of the variation in community composition (PERMANOVA *R*² = 0.034, *P* = 0.001; Supplementary Fig. S1G). Mean species relative abundances across the two groups were positively correlated (Spearman ρ = 0.696, *P* < 0.001), while only 17 of 1,357 species (1.3%) differed significantly between sources (MaAsLin3, *q* < 0.05). These included higher abundances of *Eubacterium rectale* and *Ruminococcus callidus* in VinCAPR participants and of *Lawsonibacter asaccharolyticus* in newly recruited participants (Supplementary Fig. S1H).

**Figure 1.**
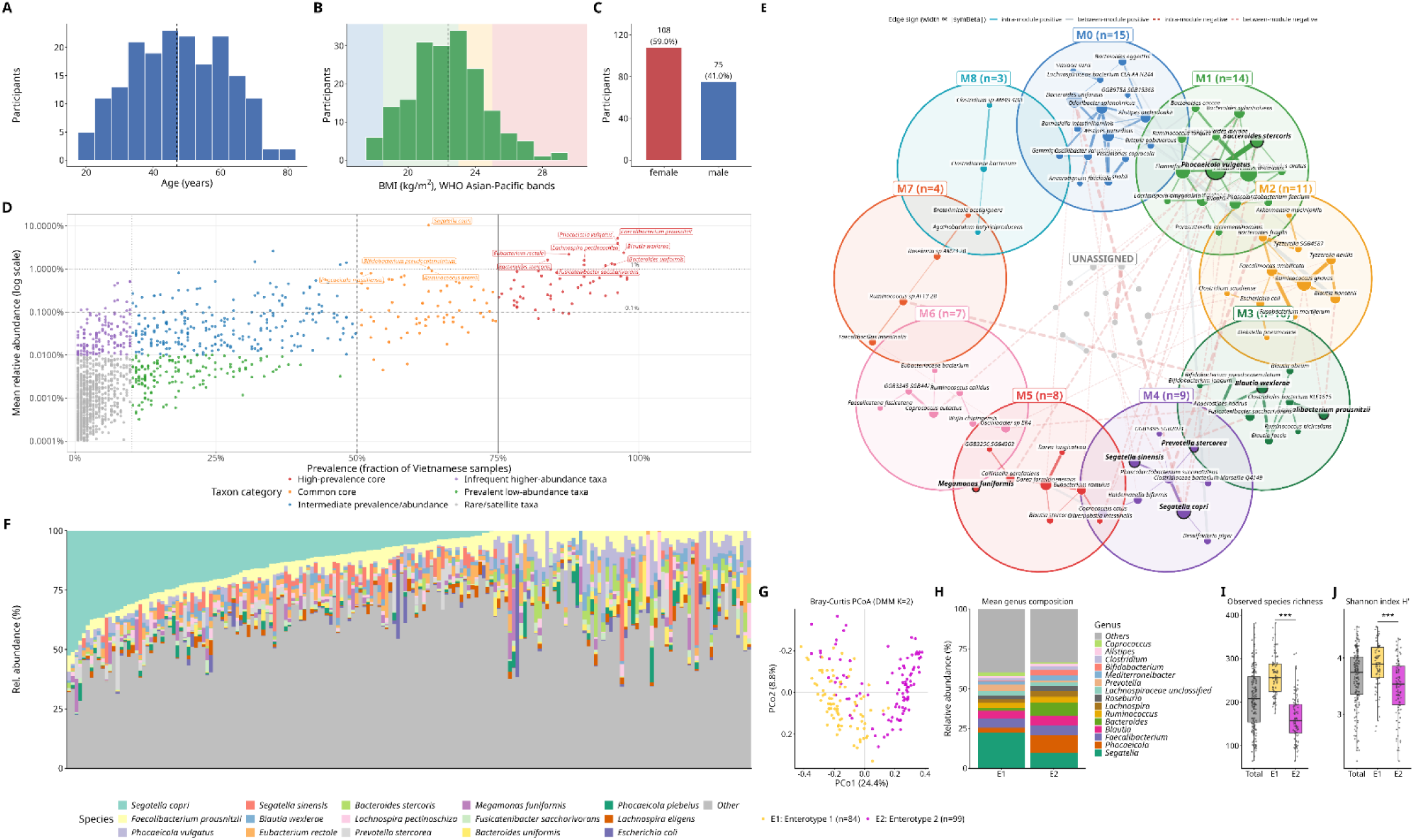
Participant characteristics and gut microbial community structure in the urban Vietnamese cohort. (A–C) Age, BMI and sex distributions among 183 participants. Vertical dashed lines in A and B indicate medians. (D) Prevalence and mean relative abundance of species-level genome bins; selected common or abundant taxa are labeled. (E) Species association network inferred using SPIEC-EASI, retaining species with prevalence ≥20% and mean relative abundance >0.1%. Node colors indicate network modules, node size represents degree, and edge width represents the absolute association estimate. Solid and dashed edges indicate positive and negative associations, respectively. (F) Species-level relative-abundance profiles of individual participants. (G) PCoA of Bray–Curtis dissimilarities, colored by Enterotype 1 and Enterotype 2 assignments from a two-component DMM model. (H) Mean genus-level relative abundant of Enterotype 1 and 2 (I-J) Observed species richness, Shannon diversity, respectively, in the complete cohort and each enterotype. Points represent individual participants. Yellow and magenta identify Enterotype 1 and Enterotype 2, respectively, *P < 0.001 (*\*\*\*).

The Vietnamese cohort containing core species such as *Faecalibacterium prausnitzii*, *Phocaeicola vulgatus* and *Blautia wexlerae*, each had prevalence above 90% and mean relative abundance above 1% (Fig. 1D). *Segatella copri* had the highest mean relative abundance, exceeding 10%, but varied substantially among participants and contributed more consistently to Enterotype 1 (Fig. 1F; Supplementary Fig. S2D). *Segatella copri* had the highest mean relative abundance among the taxa (>10%), although its contribution varied markedly among participants and it was not universally detected (Fig. 1D, F). The co-occurrence network displayed nine species modules (Fig. 1E). Notably, the core species *F. prausnitzii* and *B.wexlerae* belong to the same module M3, while *P. vulgatus* shares the same module with other *Bacteroides* species such as *B. stercoris*. *S. copri* co-occured with other *Segatella* and closely related *Prevotella* species (*S. sinensis* and *P. stercorea*), belonging to M4.

A two-component DMM model was selected to define Enterotype 1 (E1; *n* = 84) and Enterotype 2 (E2; *n* = 99) for subsequent comparisons (Supplementary Fig. S2B). The enterotypes showed partial separation in Bray–Curtis ordination space, with overlapping distributions (Fig. 1G). Enterotype was associated with community composition in covariate-adjusted PERMANOVA (*R*² = 0.067, *P* = 0.001). E1 had a higher mean relative abundance of *Segatella*, whereas E2 had higher mean relative abundances of *Phocaeicola* and *Bacteroides* (Fig. 1H; Supplementary Fig. S2D). E1 also had greater observed species richness and Shannon diversity than E2 (both *P* < 0.001; Fig. 1I,J). Species-level analyses adjusted for age, sex, and BMI identified 203 species with higher mean relative abundance in E1 and 56 species in E2 (*q* < 0.05; Supplementary Fig. S3A). Both enterotypes retained the full-cohort grouping of *F. prausnitzii* with *B. wexlerae*, while differing in the *Segatella*- and *Bacteroides*-associated regions of the network. E1 retained association between *S. copri* and *Phascolarctobacterium succinatutens*, whereas E2 contained a larger module involving *P. vulgatus* and *Bacteroides* species (17 versus 13 taxa), indicating differences in how these components of the full-cohort network were organized within each enterotype (Fig. 1E; Supplementary Fig. S3B).

Age, BMI, and sex distributions did not differ significantly between enterotypes (Supplementary Fig. S2E–G). E1 was more frequent among VinCAPR participants than newly recruited participants (57/97, 58.8%, versus 27/86, 31.4%; *P* < 0.001; Supplementary Fig. S2H). However, after matching on age, sex, and BMI, the association between recruitment source and enterotype was no longer statistically significant (*P* = 0.447; Supplementary Fig. S2I). Of the 17 recruitment source-associated species, nine are also differed by enterotypes. Most enterotype-associated species were not identified in the source comparison (Supplementary Fig. S3A). These findings suggest that the enterotype distinction was not solely attributable to recruitment source, although residual effects of recruitment and sample handling cannot be excluded.

### Vietnamese enterotypes span global compositional variation

We compared the Vietnamese cohort with public metagenomes grouped as Africa, Europe, the United States, high-income Asia and low-to-middle income Asia (Supplementary Table 1). Global ordination showed a curved, overlapping distribution of species-level profiles, with the first two coordinates explaining 12.6% and 7.0% of variation (Fig. 2D). Enterotype 1 overlapped predominantly with African and low-to-middle-Asia samples, whereas Enterotype 2 extended toward European, high-income-Asia and United States samples (Fig. 2E).

**Figure 2.**
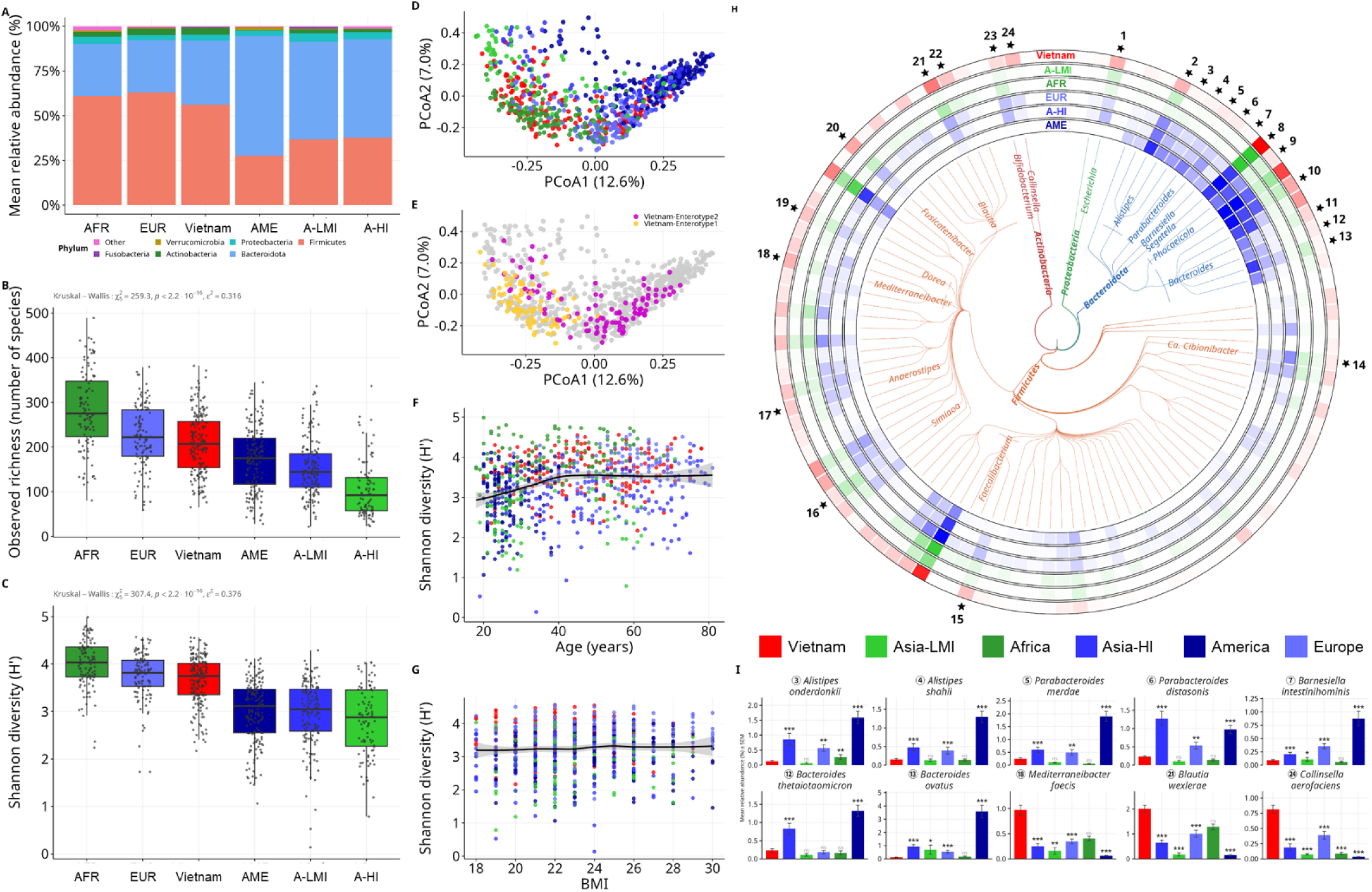
Taxonomic composition and diversity of the Vietnamese cohort in a global context. (A) Mean phylum-level relative abundance across Vietnam and five comparator groups: Africa (AFR), Europe (EUR), the United States (AME), high-income Asia (AS-d) and low-to-middle income Asia (AS-g). Comparator cohort details are provided in Supplementary Table 1. (B,C) Observed species richness and Shannon diversity across groups. Points represent individual samples; overall differences were assessed using Kruskal–Wallis tests. (D) PCoA of species-level profiles, colored by geographic group. (E) The same ordination highlighting Vietnamese participants assigned to Enterotype 1 (yellow) or Enterotype 2 (magenta); public comparator samples are gray. (F,G) Shannon diversity in relation to age and BMI, respectively. Black curves show fitted trends, with gray uncertainty bands. (H) Phylogenetic distribution and mean relative abundance of the displayed species. Branch colors indicate phylum; concentric abundance rings represent Vietnam, AS-g, AFR, EUR, AS-d and AME, from outside to inside. Color intensity indicates mean relative abundance, and numbered labels identify differentially abundant species. (I) Mean relative abundance ± standard error of the mean for ten selected species. MaAsLin3 contrasts used Vietnam as the reference and adjusted for age, sex and sequencing depth. *q* values were corrected using the Benjamini–Hochberg procedure. n.s., *q* ≥ 0.05; *q* < 0.05 (\**), <0.01 (**) and <0.001 (\*\*\**).

Observed species richness and Shannon diversity differed across geographic groups (both Kruskal–Wallis *P* < 0.001; Fig. 2B,C). Vietnamese medians were below those of the African group and above those of the Asian and United States groups, with substantial overlap with Europe. The relatively high diversity in Europe and low diversity in low-to-middle income Asia did not follow a uniform ordering by development category. Across the pooled dataset, the fitted Shannon diversity curve increased from younger ages toward mid-adulthood and then broadly leveled off, while varying little across the displayed BMI range (Fig. 2F,G).

Firmicutes and Bacteroidota dominated mean phylum-level profiles, but species-level differences extended across multiple lineages (Fig. 2A,H; Supplementary Fig. S3A). After adjustment for age, sex and sequencing depth, all seven Bacteroidota species highlighted in Fig. 2I had lower relative abundance in Vietnam than in high-income Asia and the United States, including *Bacteroides ovatus* and *B. thetaiotaomicron* (MaAslin3 adjusted *q* < 0.001). Conversely, *Mediterraneibacter faecis* and *Blautia wexlerae* had higher relative abundance in Vietnam than in high-income Asia, Europe and the United States (*q* < 0.001), but did not differ significantly from the African group. *Collinsella aerofaciens* had higher relative abundance in Vietnam than in every comparator group (*q* < 0.001). The direction for the two *Bacteroides* species is consistent at genus level with an observational comparison reporting greater *Bacteroides* relative abundance in a US urban community than in traditional Peruvian communities^33^.

A random forest classifier distinguishing Vietnamese samples from public comparator samples yielded an area under the receiver operating characteristic curve (AUC) of 0.983 (Supplementary Fig. S4B). The classifier was restricted to core taxa, demonstrating that discrimination was supported by relatively prevalent and abundant community members. Largest feature contributions include *Ruminococcus* sp. AF13_28, *Romboutsia timonensis*, and other *Faecalibacterium* and *Blautia* species (Supplementary Fig. S4C). This discrimination may partly reflect differences in sample collection and processing between studies and cannot be attributed solely to biological differences between populations.

### Enterotype alignment extends to encoded functional potential

To assess whether within-cohort differences corresponded to global industrialization-associated patterns, we applied species-based and pathway-based scores derived from public cohorts. Enterotype 2 occupied higher portions of both score distributions than Enterotype 1, although pathway scores overlapped substantially between enterotypes (Fig. 3C,E). Feature-level comparisons identified species and pathways with concordant global and within-Vietnam effect directions, alongside opposite-direction associations among species (Fig. 3D,F).

**Figure 3.**
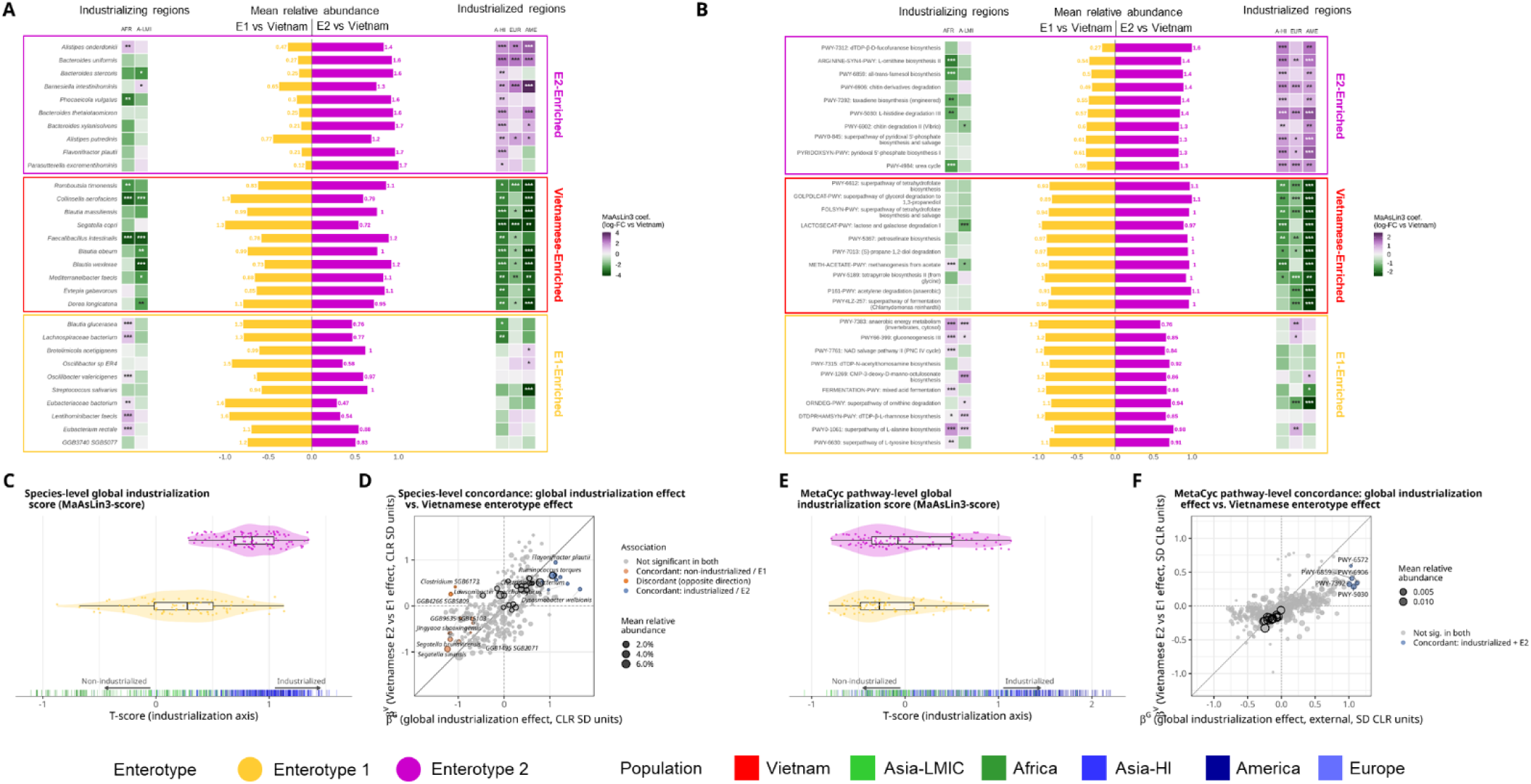
Alignment of Vietnamese enterotypes with global taxonomic and functional patterns associated with industrialization. (A,B) Selected species and MetaCyc pathways, respectively, compared across geographic groups and between Vietnamese enterotypes. Flanking heatmaps show MaAsLin3 coefficients for comparator groups relative to Vietnam; purple and green indicate positive and negative coefficients, respectively. Central mirrored bars show mean abundance ratio in Enterotype 1 (yellow, left) and Enterotype 2 (magenta, right) compared to mean across all 183 Vietnamese participants. Heatmap significance annotations indicate Benjamini–Hochberg-adjusted *q* < 0.05 (\**), <0.01 (**) and <0.001 (\*\*\**). (C,E) Distributions of species-based and pathway-based industrialization scores, respectively, for the two enterotypes. Scores were derived from coefficient-weighted MaAsLin3 models fitted to public cohorts and applied unchanged to Vietnamese samples. Scores were rescaled to reference medians of 0 for non-industrialized populations and 1 for industrialized populations. Points represent Vietnamese participants; rug marks indicate scores in public reference populations. (D,F) Concordance between global industrialization-associated effects and Vietnamese enterotype-associated effects for species and pathways, respectively. Horizontal axes show global effects; vertical axes show Enterotype 2 relative to Enterotype 1. Effects are expressed in standard-deviation units on the centered log-ratio scale. Features significant in both comparisons at Benjamini–Hochberg-adjusted *q* < 0.05 are classified as concordant or discordant according to effect direction. Point size indicates mean relative abundance. Dotted lines mark zero effects, and the diagonal marks equal effect estimates.

The taxonomic correspondence included contrasting distributions of *Bacteroides* and *Segatella*. Mean *B. uniformis* abundance was 0.27 times the Vietnamese cohort mean in Enterotype 1 and 1.6 times that mean in Enterotype 2 (Fig. 3A). Its relative abundance was also higher in high-income Asia, Europe and the United States than in Vietnam (all *q* < 0.001). *S. copri* showed the reverse pattern: mean abundance was 1.3 and 0.72 times the cohort mean in Enterotype 1 and Enterotype 2, respectively, and its relative abundance was lower in all three high-income comparator groups than in Vietnam (all *q* < 0.01). *S. brunsvicensis* and *S. sinensis* also showed concordant associations with non-industrialized reference populations and Enterotype 1 (Fig. 3D). This pattern is directionally consistent at the species-complex level with an observational analysis of human metagenomes reporting greater carriage and co-occurrence of *S. copri* complex species in non-Westernized populations^34^.

Encoded pathway profiles showed related differences. L-ornithine biosynthesis II and L-histidine degradation III each averaged 1.4 times the cohort mean in Enterotype 2, compared with 0.54 and 0.57 times the mean, respectively, in Enterotype 1 (Fig. 3B). Both pathways had higher abundance in the three high-income comparator groups than in Vietnam, with *q* < 0.01 for ornithine biosynthesis and *q* < 0.001 for histidine degradation. Mixed acid fermentation and ornithine degradation instead had higher mean abundance in Enterotype 1. The coexistence of compositional and encoded functional differences is broadly consistent with an observational study of Hmong and Karen populations spanning Thailand and the United States, whose Hmong metagenomic comparison identified differences in carbohydrate-degradation potential alongside taxonomic changes^35^

Other taxonomic and functional features were shared across the two Vietnamese configurations. *Blautia massiliensis* had mean-abundance ratios close to one in both enterotypes, despite lower relative abundance in all three high-income comparator groups than in Vietnam (all *q* < 0.05; Fig. 3A). Tetrahydrofolate biosynthesis and biosynthesis/salvage superpathways likewise had similar mean abundance between enterotypes but lower abundance in those comparator groups (all *q* < 0.01; Fig. 3B). These observations support selective correspondence between enterotype-associated taxonomic and encoded functional profiles and global population patterns, alongside Vietnamese population features shared across the two configurations.

### Enterotypes differ in carbohydrate pathways and archaeal contributions

In examining which pathway categories and species contributions distinguished the enterotypes. Enterotype 2 had greater relative abundance fractions for carbohydrate and glycan degradation and vitamin biosynthesis, whereas Enterotype 1 had a greater amino-acid-biosynthesis fraction (Fig. 4A). Individual pathways differed within these broad categories (Fig. 4B).

**Figure 4.**
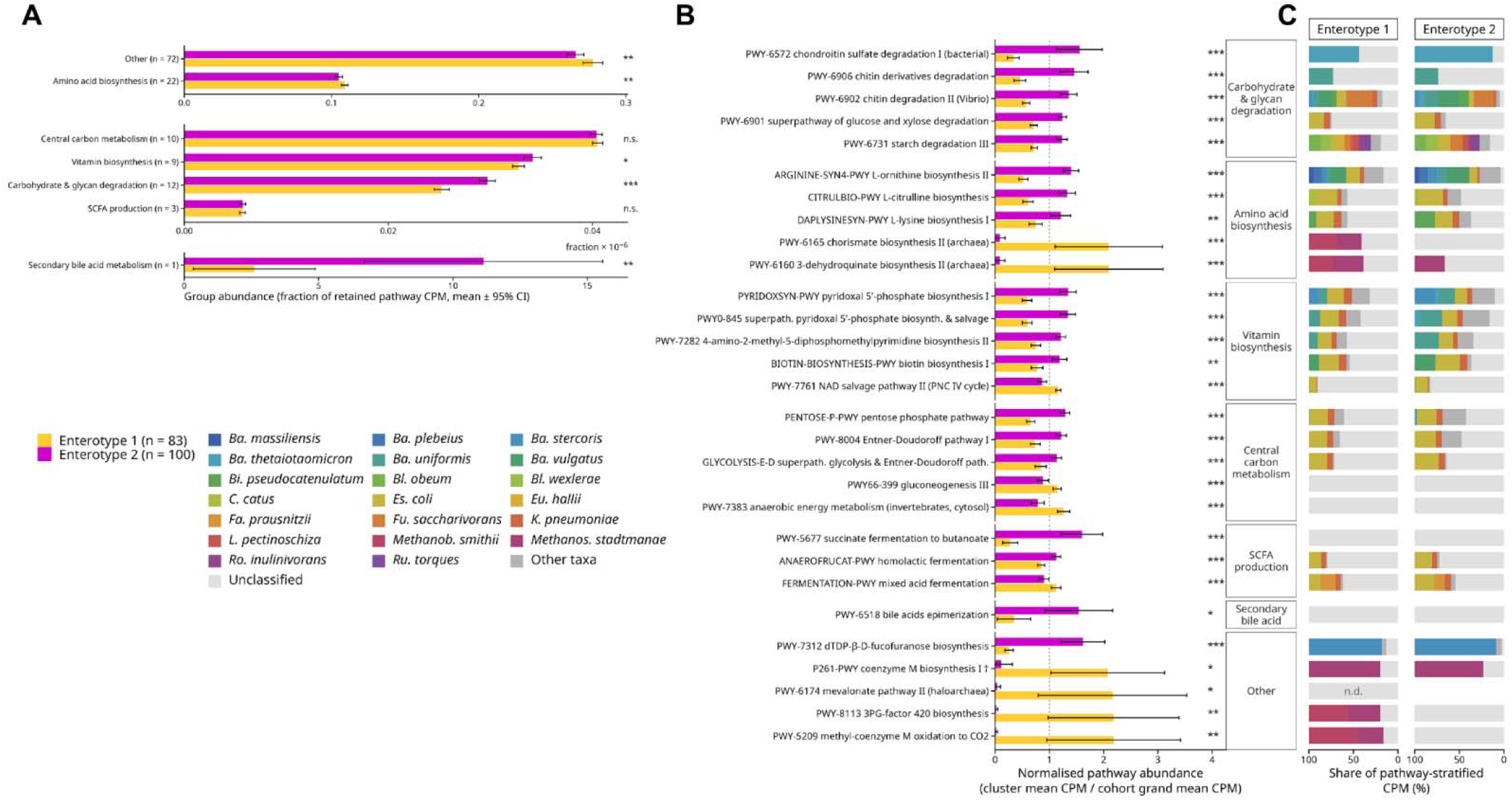
Encoded functional potential and species contributions differ between Vietnamese enterotypes. (A) Mean abundance of MetaCyc pathway groups, expressed as a fraction of the abundance of retained pathways quantified in copies per million (CPM). Intervals indicate 95% confidence intervals. Numbers in parentheses indicate the number of pathways assigned to each functional group. (B) Selected pathway abundances in Enterotype 1 and Enterotype 2, normalized to the corresponding mean pathway abundance across the Vietnamese cohort. Yellow and magenta identify Enterotype 1 and Enterotype 2, respectively, in A and B. (C) Species-stratified contributions to the pathways shown in B, estimated using HUMAnN. Mirrored stacked bars show the mean percentage of pathway abundance attributed to individual species in Enterotype 1 (left) and Enterotype 2 (right). Segment colors identify species; dark gray indicates other species and light gray indicates unclassified contributions. Pathway profiles represent encoded functional potential.

All five displayed carbohydrate-degradation pathways had greater normalized mean relative abundance in Enterotype 2, including chondroitin sulfate degradation I, chitin-derivative degradation and starch degradation III (Fig. 4B). Species-stratified profiles identified *B. thetaiotaomicron* as the principal classified contributor to chondroitin sulfate degradation, with a larger contribution in Enterotype 2 (Fig. 4C). *B. uniformis* contributed to chitin-derivative degradation, alongside a substantial unclassified fraction.

Biosynthetic pathways showed contrasting directions within functional groups. Enterotype 2 had greater normalized mean relative abundance of pyridoxal 5′-phosphate and biotin biosynthesis pathways (Fig. 4B). Enterotype 1 had greater relative abundance of archaeal chorismate and 3-dehydroquinate biosynthesis pathways, together with coenzyme M and F420 biosynthesis annotations. HUMAnN attributed classified contributions to these archaeal-associated profiles to *Methanobrevibacter smithii* and *Methanosphaera stadtmanae* (Fig. 4C). The higher aggregate amino-acid-biosynthesis fraction in Enterotype 1 coexisted with higher means for individual ornithine, citrulline and lysine biosynthesis pathways in Enterotype 2 (Fig. 4A,B).).

Aggregate central-carbon metabolism and the category annotated as “SCFA production” did not differ significantly between enterotypes (Fig. 4A). Their component pathways nevertheless varied, i.e., succinate fermentation to butanoate and homolactic fermentation had higher normalized mean relative abundance in Enterotype 2, whereas mixed acid fermentation had a higher mean in Enterotype 1 (Fig. 4B). Bile-acid epimerization also had a higher mean in Enterotype 2, although its species contributions were unclassified (Fig. 4B,C).

### Lifestyle variables moderately distinguish enterotypes

Lifestyle data were available for 105 participants (Supplementary Fig. S1A). Comsumption of alcohol (weekly) and nuts/peanuts in diet were associated with higher odds of Enterotype 1 after adjustment for age, sex and BMI and correction across the 12 tested variables (both adjusted *P* < 0.05; Fig. 5A). The displayed associations with overall diet score and walking duration did not meet the corrected significance threshold.

**Figure 5.**
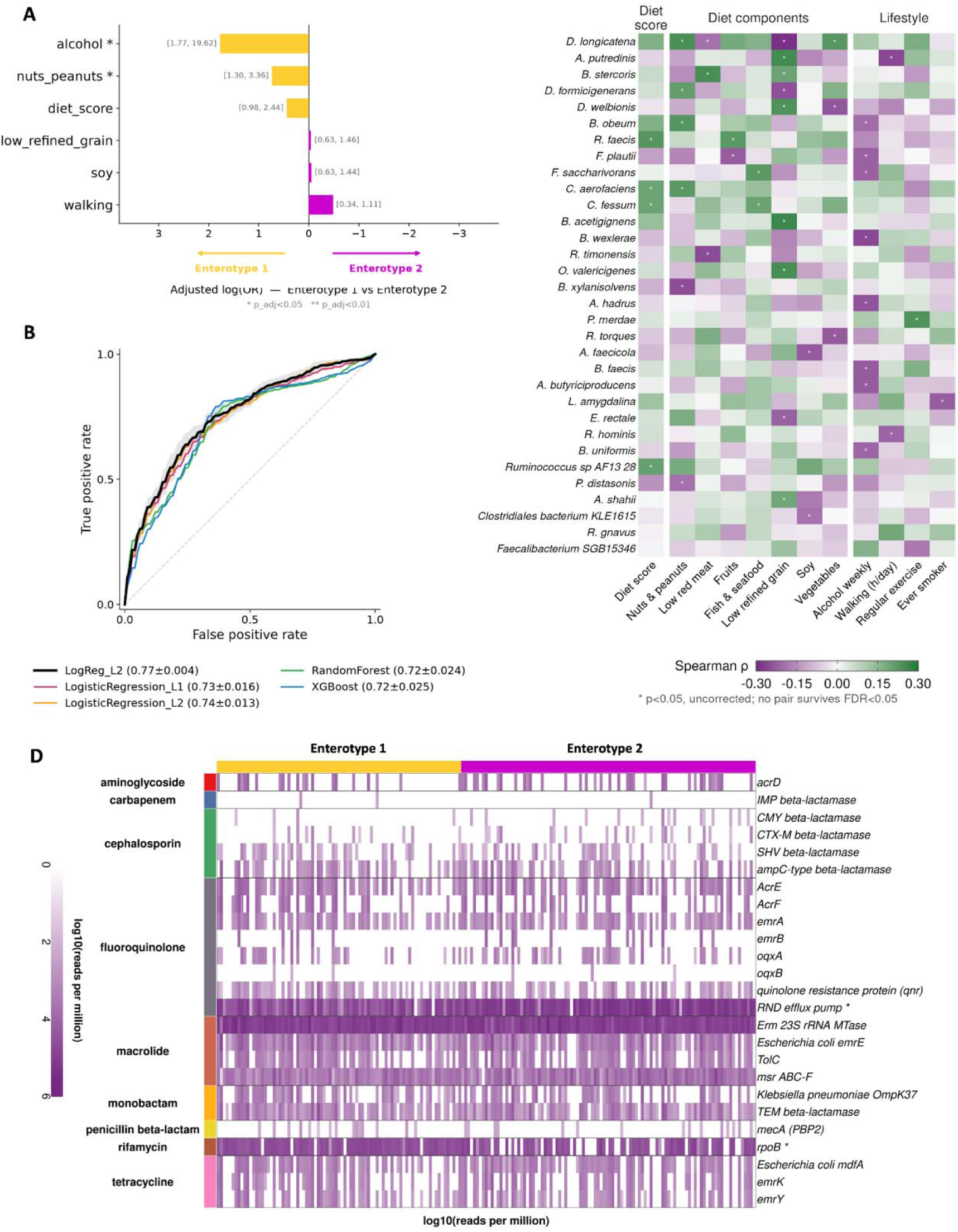
Lifestyle associations and antimicrobial-resistance gene profiles of Vietnamese enterotypes. (A) Associations between dietary or lifestyle variables and Enterotype 1 relative to Enterotype 2, estimated using separate logistic regression models adjusted for age, sex and BMI. The horizontal axis shows log odds ratios; bracketed values indicate odds-ratio intervals. Positive and negative estimates indicate associations with Enterotype 1 and Enterotype 2, respectively. *P* values were corrected across 12 tests using the Benjamini–Hochberg procedure. (B) Receiver operating characteristic curves for enterotype classification among participants with complete lifestyle data. Models include an L2-penalized logistic classifier using selected variables and classifiers incorporating all 12 variables: L1- and L2-penalized logistic regression, random forest and XGBoost. Models were evaluated using repeated stratified five-fold cross-validation with 20 repeats. Legend values report mean ± standard deviation of the area under the curve (AUC) across repeats. (C) Spearman correlations between selected species and dietary or lifestyle variables. Green and purple indicate positive and negative correlations, respectively. No association remained significant after multiple-testing correction. (D) ARG profiles across individual participants, grouped by enterotype. Rows represent resistance-associated genes or gene families organized into nine antibiotic classes; color intensity indicates log-transformed RPM. Asterisks identify gene-wise enterotype differences from Wilcoxon rank-sum tests with Benjamini–Hochberg-adjusted P. *P* < 0.05. *P* < 0.05 (*).

An L2-penalized logistic classifier using selected lifestyle variables (alcohol, nuts, walking) achieved a mean cross-validated AUC of 0.77 ± 0.004 (mean ± standard deviation; Fig. 5B). Models incorporating all 12 variables yielded mean AUCs of 0.72–0.74 across penalized logistic regression, random forest and XGBoost. These models were evaluated using repeated stratified five-fold cross-validation with 20 repeats.

Exploratory species-level analyses identified nominal associations with these exposures. The nuts-and-peanuts measure correlated positively with *Dorea longicatena* and *Blautia obeum*, whereas weekly alcohol use correlated negatively with *Anaerostipes hadrus* and *B. uniformis* (nominal *P* < 0.05; Fig. 5C). No species–exposure association remained significant after false-discovery-rate correction at 0.05.

### Antimicrobial resistance-associated profiles are largely shared across enterotypes

Antimicrobial resistance-associated profiles were heterogeneous within both enterotypes, with broadly distributed Erm 23S rRNA methyltransferase and msr ABC-F annotations (Fig. 5D). Among the β-lactamase families examined at variant level, SHV annotations were more widespread across the cohort, whereas CTX-M and CMY annotations occurred in smaller subsets of participants and IMP annotations were rare (Fig. 5D; Supplementary Fig. S5).

Both enterotypes carried several of the same β-lactamase variants. A wider range of CTX-M and CMY variants was detected in E2, whereas the common SHV variants showed numerically higher prevalence in E1 (Supplementary Fig. S5B–D). These descriptive differences occurred within broadly shared resistance-associated profiles. Among the features displayed in Fig. 5D, RND efflux-pump and *rpoB* annotations differed significantly in normalized read abundance between enterotypes (Wilcoxon rank-sum tests, Benjamini–Hochberg-adjusted *P* < 0.05).

## Discussion

The central finding presented here is that microbiome differences commonly associated with global industrialization are also present within a clinically screened cohort from Hanoi. Enterotype 1 and Enterotype 2 combined contrasting taxonomic profiles with differences in encoded functional potential, extending observations from geographic and migration studies to variation within this population^33,35,36^. Both configurations also shared features that distinguished the cohort from several global comparators. A single Vietnamese population average would conceal this combination of globally aligned variation and a shared local background.

The greater richness and diversity of Enterotype 1 are consistent with an observational survey of Asian schoolchildren, in which *Prevotella*-rich communities contained more phylotypes than *Bifidobacterium/Bacteroides*-rich communities^37^. Differences in age, study design and taxonomic resolution preclude direct equivalence between these classifications. Nevertheless, both studies associate community composition with richness within Asian populations. Distinguishing between these shared and variable features provides critical biological context. Taxonomically, the enterotype differences extend beyond a simple *Prevotella*-*Bacteroides* dichotomy, situating variations within broader modules of co-varying organisms. For instance, *S. copri* clustered within an association module containing other *Segatella* and *Prevotella* species, while *P. vulgatus* grouped with several *Bacteroides* species. This is consistent with the co-occurrence of multiple members of the *S. copri* complex in non-urbanized populations, along with their distinct genetic and polysaccharide-utilization repertoires^34,38–40^. However, co-abundance associations alone do not establish direct microbial interactions or cross-feeding.

Despite these enterotype-driven variations, certain shared features underscore a distinct local baseline. For example, *B. massiliensis* and folate-related pathways maintained similar abundances across both enterotypes yet differed significantly from high-income reference populations. This conserved core microbiome provides plausible links to host physiology and establishes the cohort as a highly relevant baseline comparator for inflammatory and neoplastic gastrointestinal diseases in Vietnam. Key core members include *B. wexlerae*—which has been inversely associated with obesity and type 2 diabetes, and reduces diet-induced metabolic disease in mice^41,42^, though its acetate production reflects sugar fermentation rather than hydrogenotrophic acetogenesis^43^. The cohort is also enriched in butyrate producers with anti-inflammatory potential, such as *Fusicatenibacter saccharivorans* (Clostridium subcluster XIVa^44^), which induces interleukin-10 and ameliorates colitis in experimental systems but is depleted in active ulcerative colitis^45^, and *Mediterraneibacter faecis*, which is reduced in Crohn’s disease^46^. The high relative abundance of *Collinsella aerofaciens* further illustrates the importance of local reference data. This species has been associated with obesity and steatohepatitis in other populations, and experimental studies have reported increased circulating ethanol in mice and production of an immunologically active lipid^47,48,49^. Because microbial background distributions vary geographically, locally matched comparisons are needed to distinguish population-associated variation from disease-associated changes^50^. Without this local context, future Vietnamese case-control studies risk misclassifying a population norm as disease-associated.

Functional profiles supported the same broad alignment with global industrialization-associated variation as the taxonomic profiles. Enterotype-associated pathway differences resembled contrasts among global comparator groups, although individual species did not consistently follow the same ordering. This pattern points to functional redundancy, whereby different populations can retain comparable metabolic potential through different organisms^51^, a phenomenon similarly observed following migration from Thailand to the United States^35,52^.

Vietnam’s rice-centered food culture may be one factor that helps interpret these functional shifts. Foods eaten alongside a shared staple diversify the substrates reaching the colon; for example, nuts supply plant cell-wall polysaccharides that favor specific carbohydrate-utilizing organisms, aligning with the strain-specific utilization of these substrates by *S. copri* ^53^. The association between nut intake and Enterotype 1 may therefore reflect variation in substrate composition within a broadly shared dietary pattern. A Vietnamese peanut intervention reported selective microbiome changes without significant changes in overall alpha or beta diversity, providing experimental context for food-specific responses^54^. These observations support further investigation of dietary exposures but do not establish that they caused the community differences observed here.

These functional findings reveal biochemical contrasts, for example, while Enterotype 2 exhibited lower overall species richness, it harbored a greater relative abundance of specific carbohydrate-degradation pathways spanning chemically diverse substrates, including starch, chitin derivatives, and chondroitin sulfate (the latter agreeing with the characterized enzymatic capacity of *B. thetaiotaomicron*^55^). Therefore, microbial utilization of staple starch versus the diverse polysaccharides from accompanying foods drives distinct community repertoires. Similar substrate-specific nuance was evident in amino-acid metabolism, whereby Enterotype 1 featured a higher aggregate biosynthesis fraction, whereas several individual biosynthetic pathways were elevated in Enterotype 2. Opposing patterns of ornithine biosynthesis and degradation further illustrate the differing potential of these enterotypes to synthesize versus consume the same nitrogen-containing compound.

Methanogen-associated pathways enriched in Enterotype 1 were largely attributable to *Methanobrevibacter smithii* and *Methanosphaera stadtmanae*, organisms with distinct methanogenesis-related gene repertoires^56^. Experimental evidence that methanogens consume hydrogen and influence bacterial fermentation provides a rationale for investigating their ecological roles^57^. The two species may also differ in host relevance, for example, *M. stadtmanae* elicits stronger pro-inflammatory cytokine responses than *M. smithii*^58^, while archaeal composition varies with geography, urbanization, and diet^59^. However, hydrogen consumption, methane production and fermentation rates were not measured here. The present results therefore only identify differences in methanogen-associated gene content.

The resistome, by contrast, was broadly shared across the two Vietnamese enterotypes, despite their taxonomic and functional differences. Clinically important β-lactamase genes, including *blaCTX-M* and *blaCMY*, warrant particular attention because of their relevance to third-generation cephalosporin resistance in Enterobacterales, a WHO critical-priority category. This distinction is important in Vietnam, where healthy people harbor a substantial reservoir of resistance genes and patterns of antibiotic use differ from those in many surveyed populations^17,60^. Exclusion of recent antibiotic use reduces, but does not remove, treatment-related effects because resistome changes may persist for months^61,62^. Antimicrobial exposure, community structure, environmental transmission and mobile elements can all shape gut resistomes^63,64^; read-based data cannot separate these mechanisms or demonstrate horizontal transfer. Genome-resolved linkage to microbial hosts and mobile elements will be required^65^.

The cohort’s clinical screening strengthens its value as a reference, but several limitations still remain. First, First, geographic representation is limited: recruitment was concentrated in Hanoi, limiting generalizability to the wider Vietnamese population. The public comparator datasets also represent a selected subset of global populations and may not capture their full microbiome diversity. Second, differences in sample handling and sequencing across the contributing datasets may influence the observed microbiome profiles. Consistent bioinformatic processing cannot fully remove technical heterogeneity between our cohort and global comparators or among public datasets^68,69^. Third, the cross-sectional design and analytical scope limit biological interpretation. The enterotypes summarize overlapping variation at a chosen clustering resolution^66,67^, and the data has not yet established progression from Enterotype 1 to Enterotype 2. Absolute microbial quantification was not performed, and strain-specific variation remains unresolved.

A key unanswered question is whether these coexisting microbiome configurations are durable features of the gut ecosystem or are modified by changes in individual exposures over time. Repeated sampling linked to dietary assessment and metabolite measurements would test that relationship while examining the specific substrate-utilization and hydrogen-exchange processes suggested here. Such evidence would extend the study’s central observation beyond description, linking the coexistence of globally associated microbiome configurations within an urban population to the ecological processes that drive their stability or change over time.

## Data Availability Statement

Data were deposited under BioProject PRJNA1446191

## Ethical Statement

All authors are accountable for all aspects of the work in ensuring that questions related to the accuracy or integrity of any part of the work are appropriately investigated and resolved.

## Funding

This work was supported by Vinmec-VinUni Institute of Immunology under Grant 23.11.SRI awarded to Xuan-Hung Nguyen.

## Financial Disclosure

The authors have no financial relationships relevant to this article to disclose.

## Conflict of Interest

The authors have no other conflicts of interest to declare.

## Author contributions

Thanh Thien Le: Conceptualization, Methodology, Investigation, Data curation, Administration, Writing - Original Draft, Writing - Review & Editing, Visualization; Joshua Tran: Validation, Formal analysis, Writing - Original Draft, Writing - Review & Editing; Linh Hong Thi Mai: Writing - Original Draft, Writing – Review & Editing; Dao Bich Thi Tran: Investigation, Data Curation, Writing - Review & editing; Dat Quoc Nguyen: Investigation, Data Curation, Writing - Review & editing; Quynh Mai Nguyen: Formal analysis, Visualization, Writing - Review & editing; Giang Minh Vu: Formal analysis, Visualization, Writing - Original Draft, Writing - Review & editing; Minh Ngoc Truong: Formal analysis, Data curation, Visualization, Writing - Review & editing Manh Khanh Dinh: Investigation, Writing - Review & editing; Hao Chung The: Writing - Review & editing; Thanh Hong Nguyen: Investigation, Writing - Review & editing; Chi Du Thi Tran: Data Curation, Writing - Review & editing; Huan Duc Pham: Investigation, Writing - Review & editing; Huong Thu Thi Pham: Investigation, Writing - Review & editing; Muoi Xuan Nguyen: Investigation, Writing - Review & editing; Thuy Xuan Nguyen: Investigation, Writing - Review & editing; Hang Dao Viet: Writing - Review & editing; May Ngoc Le: Writing - Review & editing; Tuan Thanh Nguyen: Writing - Review & editing; Jennifer Le: Writing - Review & editing; Jack Gilbert: Writing - Review & editing; Xuan-Hung Nguyen: Funding acquisition, Conceptualization, Supervision, Writing - review & editing;

## Supplementary information

**Supplementary Table 1:**
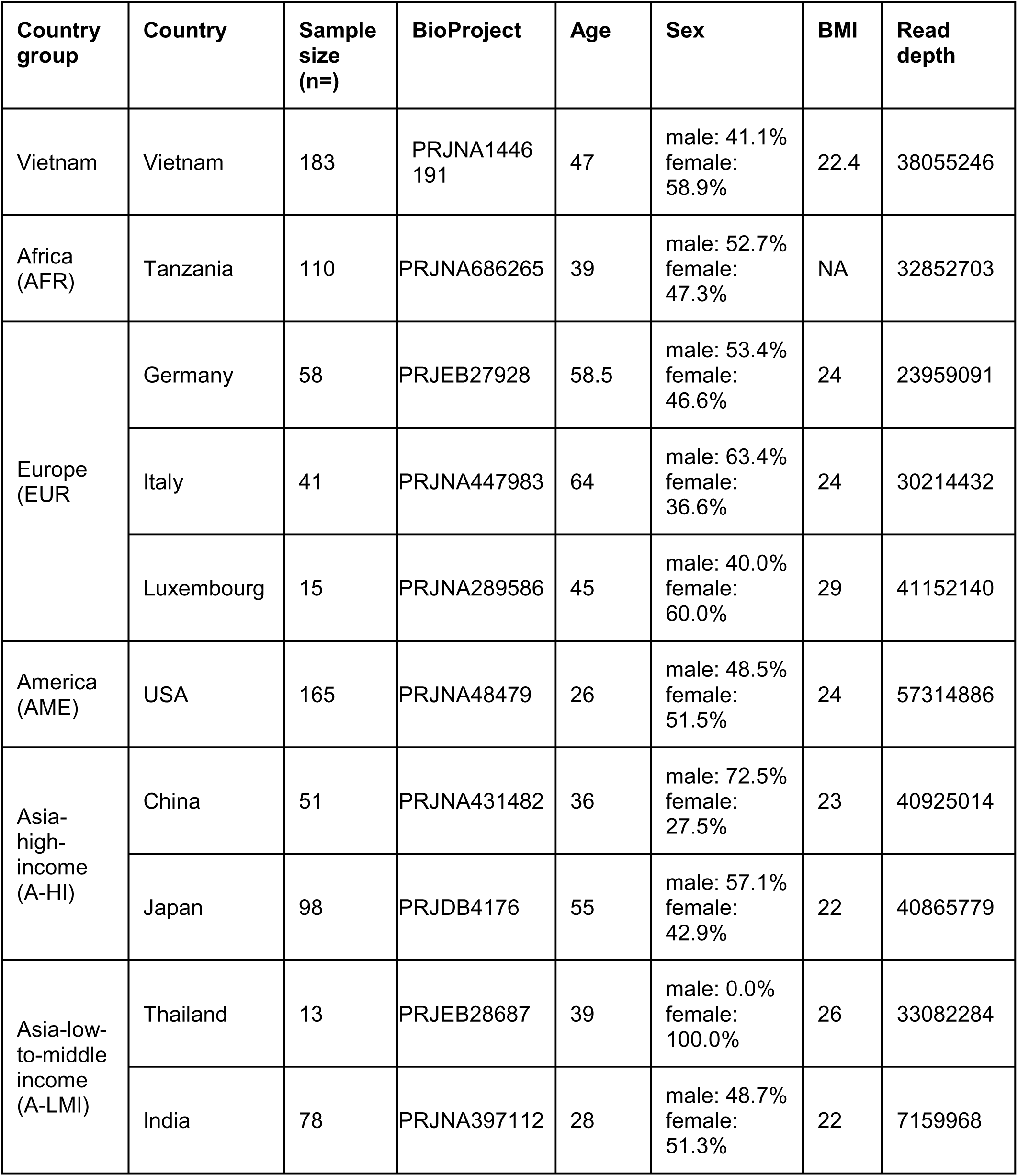
Sample characteristics.

**Supplementary Figure S1.**
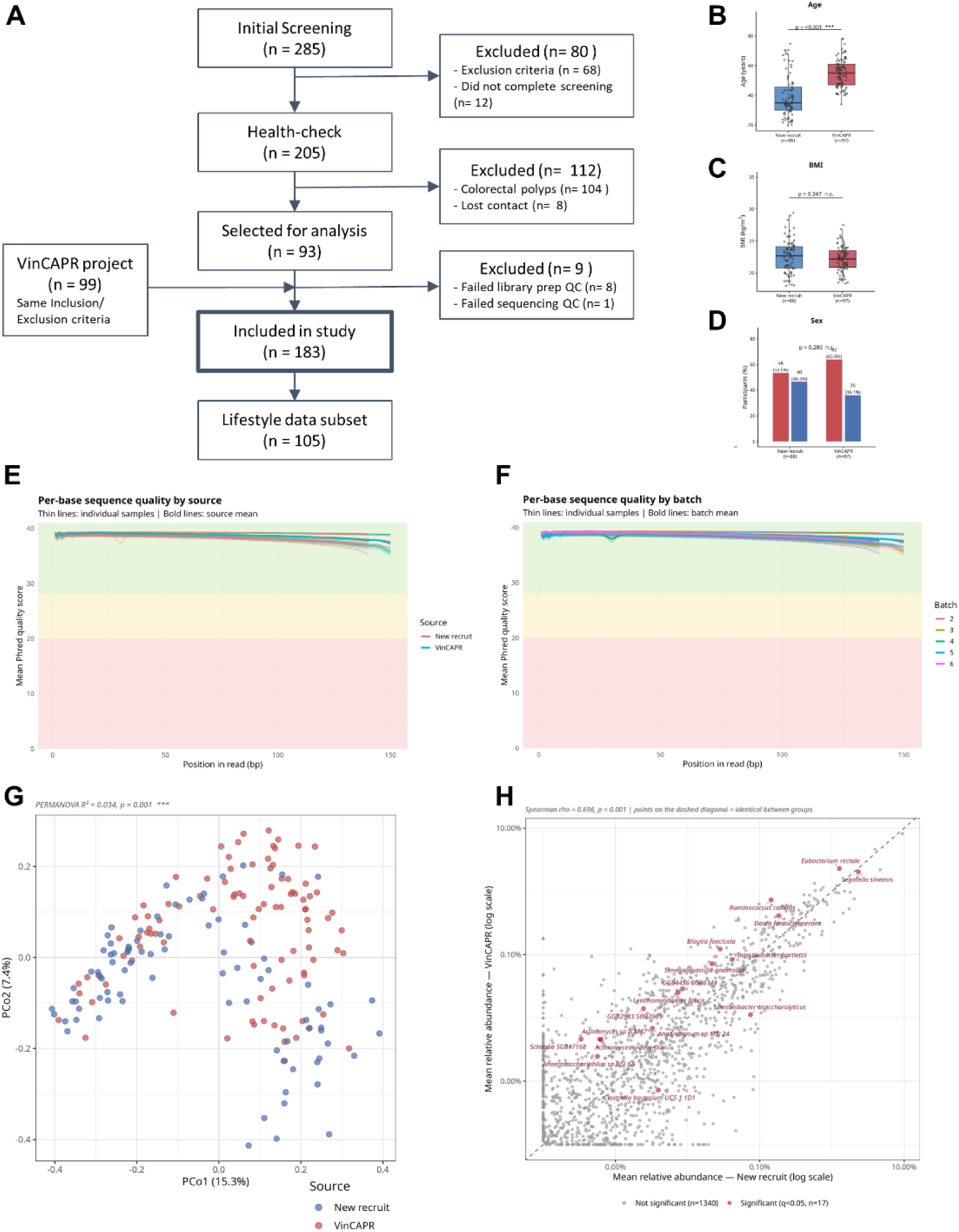
Participant selection, sequencing quality and comparisons between recruitment sources. (A) Participant screening, sample processing and inclusion in the final cohort. (B–D) Age, BMI and sex distributions, respectively, among retained participants from the newly recruited group (*n* = 86) and VinCAPR (*n* = 97). (E,F) Per-base sequencing quality by recruitment source and sequencing batch, respectively. Thin curves represent individual samples and bold curves indicate group means. (G) Principal coordinates analysis of microbial community profiles, colored by recruitment source. (H) Comparison of mean species relative abundances between newly recruited participants and VinCAPR participants on logarithmic axes. Each point represents a species; the dashed diagonal indicates equal mean relative abundance. The Spearman correlation is shown, and highlighted species have *q* < 0.05.

**Supplementary Figure S2.**
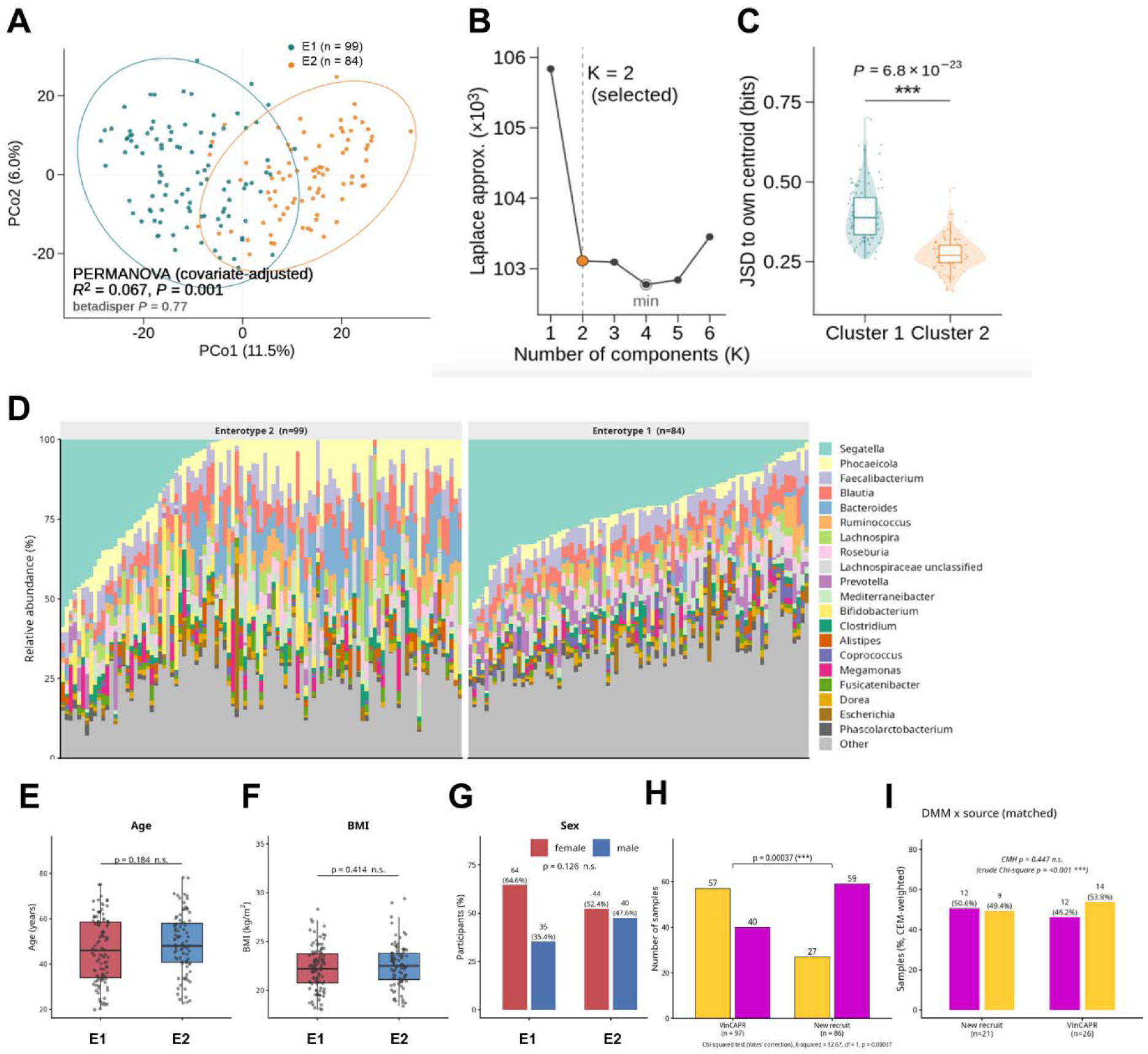
Enterotype clustering diagnostics and participant characteristics. (A) Principal coordinates analysis colored by DMM assignment. Results from PERMANOVA and the dispersion test are displayed. (B) Laplace criterion for DMM models containing one to six components. The selected two-component model and the minimum Laplace value are indicated. (C) Jensen–Shannon divergence between each sample and its assigned cluster centroid, expressed in bits. (D) Genus-level relative-abundance profiles, grouped by enterotype and ordered within groups by dominant genus. Colors identify the displayed genera; remaining genera are grouped as “Other.” (E–G) Age, body BMI and sex distributions, respectively, by enterotype. Points in E and F represent individual participants; bars in G show sex composition. (H) Enterotype counts by recruitment source (I) Enterotype distributions by recruitment source after coarsened exact matching (CEM) on age, sex, and BMI. Bar heights represent CEM-weighted percentages. n.s., not significant; ***, *P* < 0.001

**Supplementary Figure S3.**
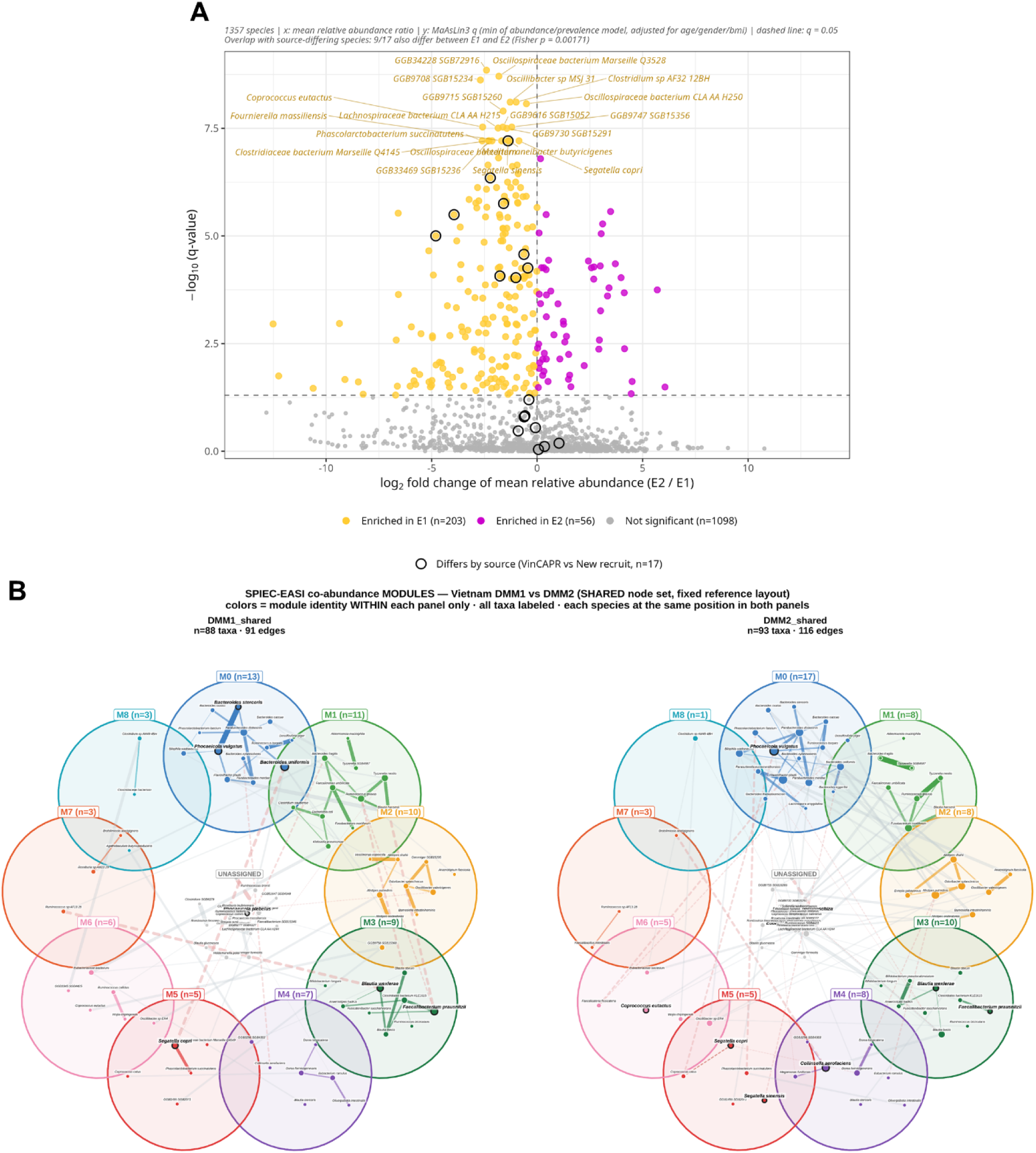
Species-level taxonomic differences and co-abundance networks by enterotype. (A) Volcano plot of associations between species and enterotype. Yellow and magenta indicate species enriched in E1 (*n* = 203) and E2 (*n* = 56), respectively; gray indicates nonsignificant species (*n* = 1,098). Dashed lines indicate equal mean relative abundance and *q* = 0.05. Black outlines identify the 17 species that differed by recruitment source, nine of which also differed between enterotypes (Fisher’s exact test, *P* = 0.00171). (B) Species co-abundance networks inferred using SPIEC-EASI for the two enterotypes. Nodes represent species, and edges represent inferred associations. Corresponding species occupy the same positions in both networks to facilitate comparison.

**Supplementary Figure S4.**
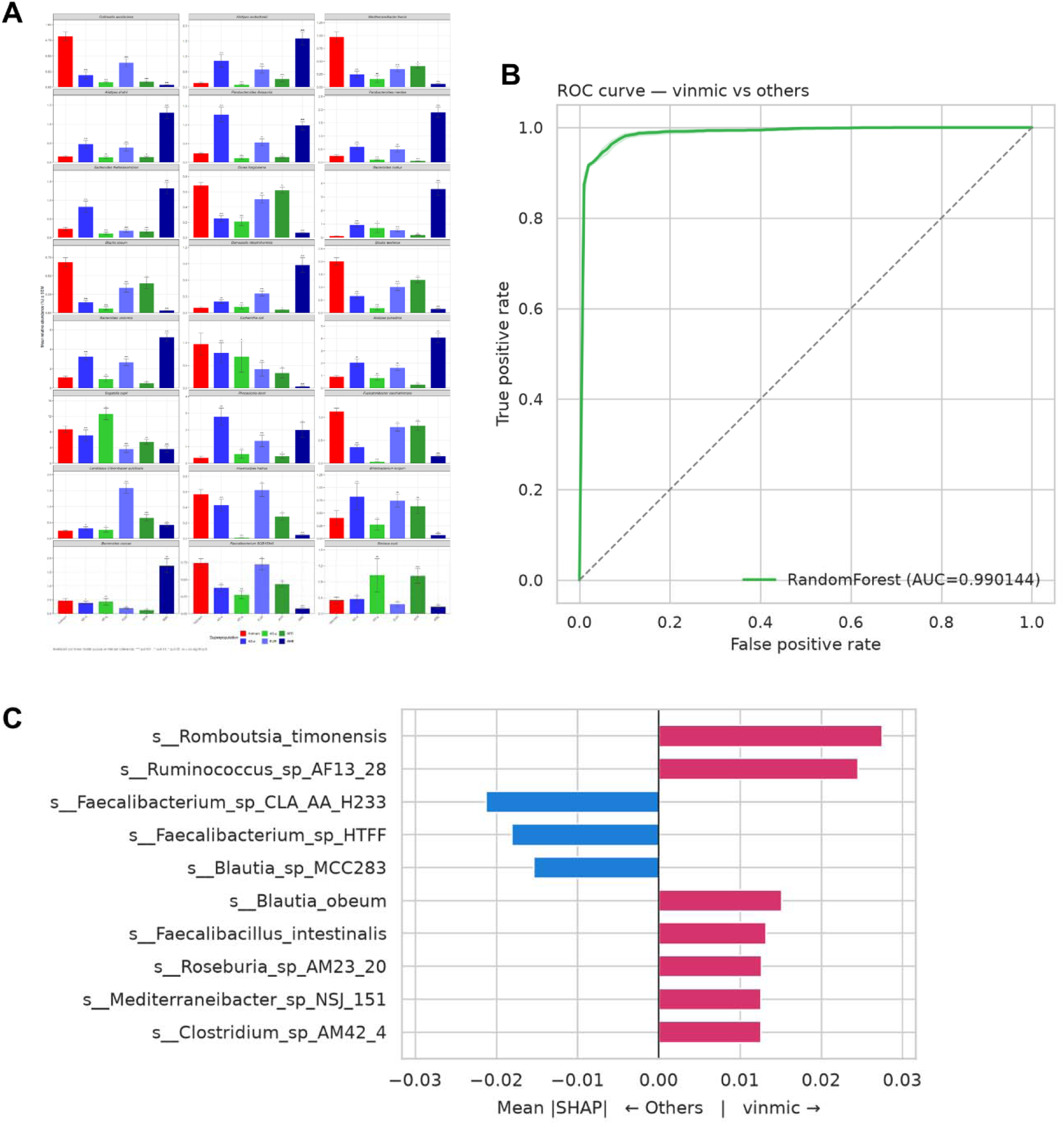
Taxonomic features distinguishing the Vietnamese cohort from public comparator populations. (A) Relative abundances of 24 selected species across Vietnam and public comparator groups. MaAsLin3 contrasts used Vietnam as the reference and adjusted for age, sex and sequencing depth. Significance annotations indicate Benjamini–Hochberg-adjusted *q* < 0.05 (\**), <0.01 (**) and <0.001 (\*\*\**). (B) Receiver operating characteristic curve for a random forest classifier distinguishing Vietnamese samples from public comparator samples using core taxa (prevalence ≥20% and mean relative abundance >0.1%) (C) SHAP-based feature contributions for ten selected species, oriented toward the Vietnamese or comparator class as indicated.

**Supplementary Figure S5.**
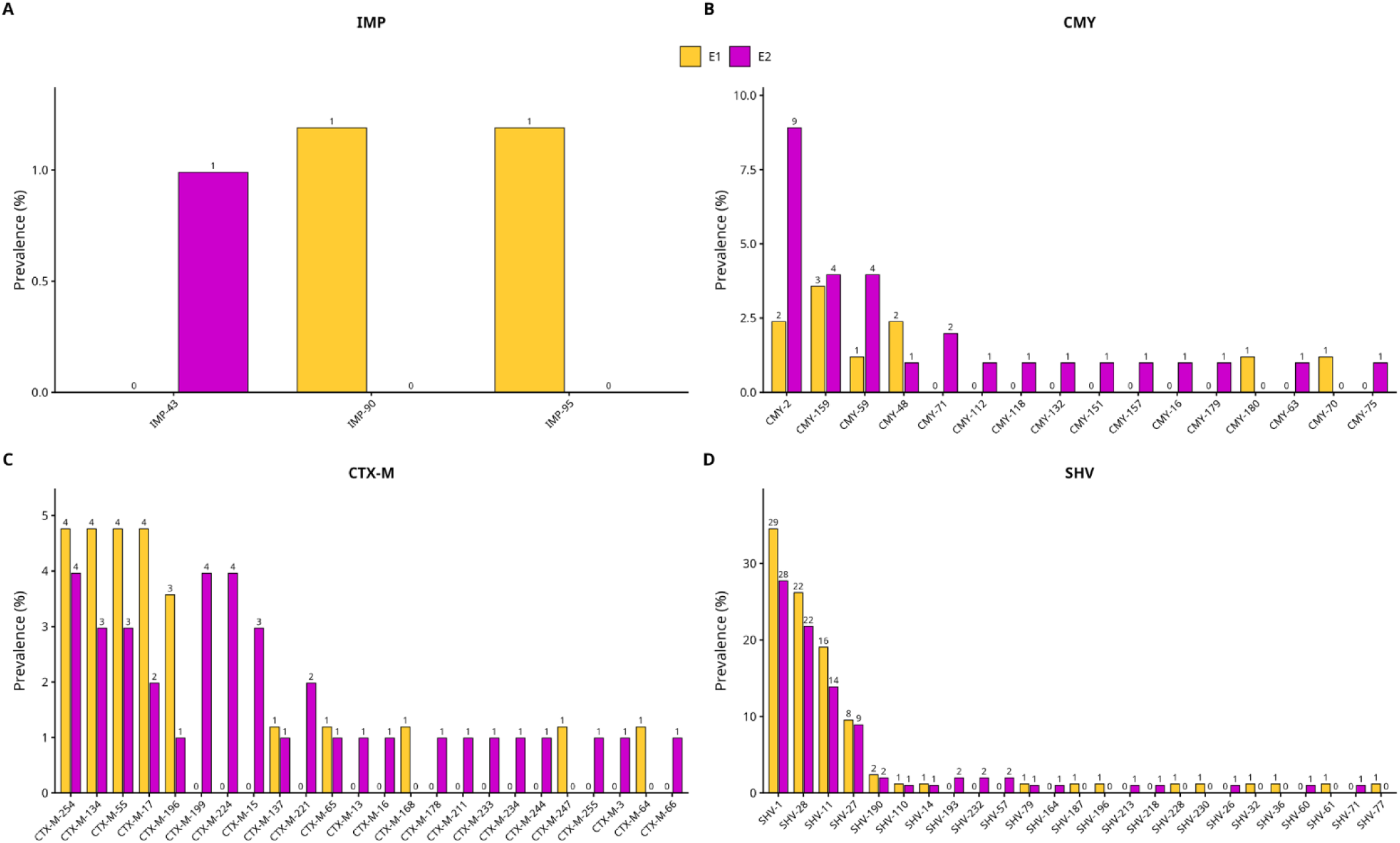
Prevalence of β-lactamase gene variants by enterotype. Prevalence of annotated variants within the (A) IMP, (B) CMY, (C) CTX-M, and (D) SHV families in Enterotype 1 (E1; *n* = 84) and Enterotype 2 (E2; *n* = 99). Bar heights indicate the percentage of samples in each enterotype with the corresponding variant; numbers above bars indicate positive-sample counts. Yellow denotes E1 and magenta denotes E2. Comparisons are descriptive; no statistical significance is indicated.

## Notes

### Competing Interest Statement

The authors have declared no competing interest.

