## Supplementary Information for "Urban Vietnam at the microbiome transition: Co-existing enterotypes reflect the global industrialization patterns"

**Supplementary Table 1: Sample characteristics**

| **Country group** | **Country** | **Sample size (n=)** | **BioProject** | **Age** | **Sex** | **BMI** | **Read depth** |
| --- | --- | --- | --- | --- | --- | --- | --- |
| Vietnam | Vietnam | 183 | PRJNA1446191 | 47 | male: 41.1% female: 58.9% | 22.4 | 38055246 |
| Africa (AFR) | Tanzania | 110 | PRJNA686265 | 39 | male: 52.7% female: 47.3% | NA | 32852703 |
| Europe  (EUR | Germany | 58 | PRJEB27928 | 58.5 | male: 53.4% female: 46.6% | 24 | 23959091 |
|  | Italy | 41 | PRJNA447983 | 64 | male: 63.4% female: 36.6% | 24 | 30214432 |
|  | Luxembourg | 15 | PRJNA289586 | 45 | male: 40.0% female: 60.0% | 29 | 41152140 |
| America  (AME) | USA | 165 | PRJNA48479 | 26 | male: 48.5% female: 51.5% | 24 | 57314886 |
| Asia-high-income  (A-HI) | China | 51 | PRJNA431482 | 36 | male: 72.5% female: 27.5% | 23 | 40925014 |
|  | Japan | 98 | PRJDB4176 | 55 | male: 57.1% female: 42.9% | 22 | 40865779 |
| Asia-low-to-middle income  (A-LMI) | Thailand | 13 | PRJEB28687 | 39 | male: 0.0% female: 100.0% | 26 | 33082284 |
|  | India | 78 | PRJNA397112 | 28 | male: 48.7% female: 51.3% | 22 | 7159968 |

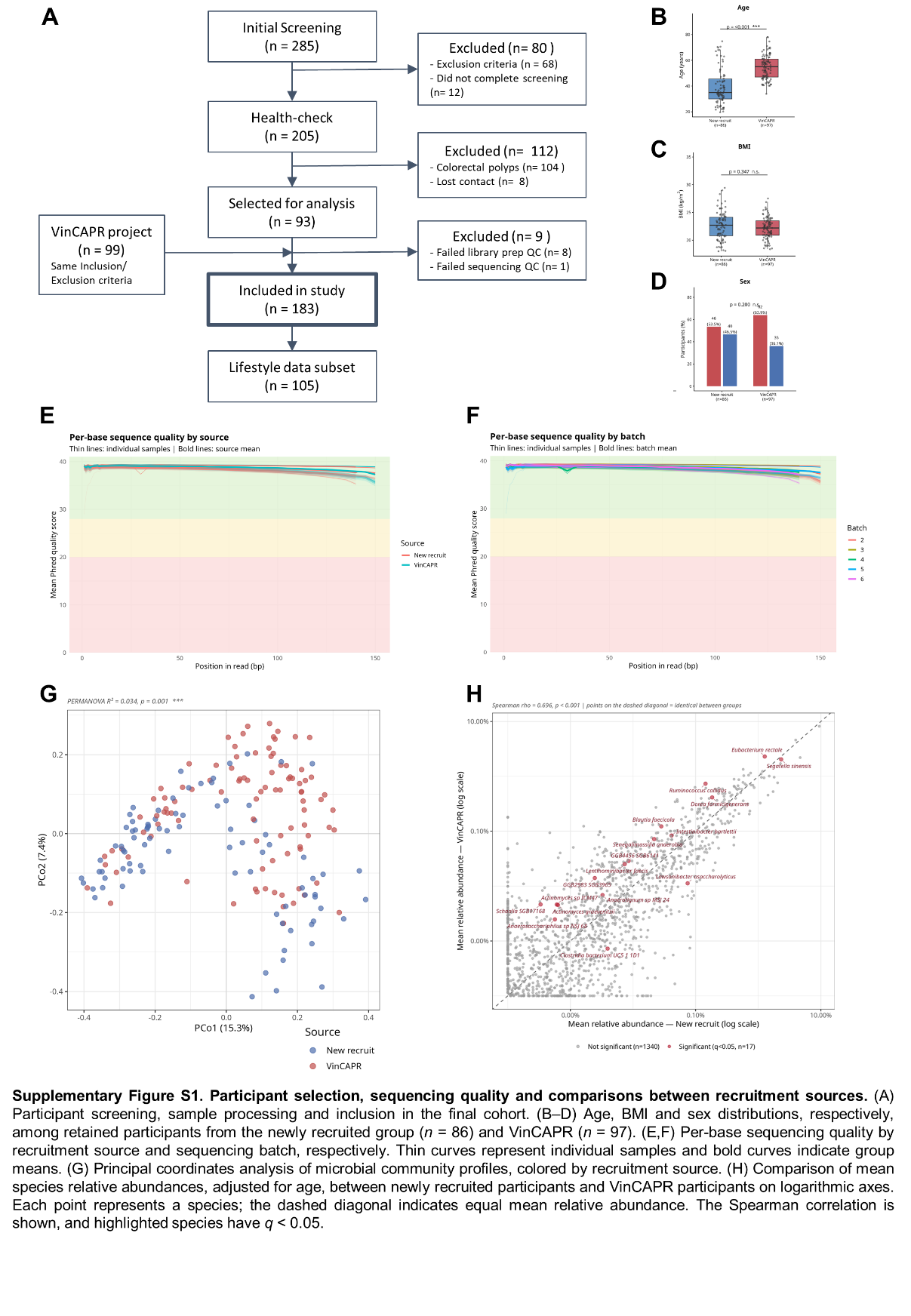

**Supplementary Figure S1. Participant selection, sequencing quality and comparisons between recruitment sources.** (A) Participant screening, sample processing and inclusion in the final cohort. (B–D) Age, BMI and sex distributions, respectively, among retained participants from the newly recruited group (*n* = 86) and VinCAPR (*n* = 97). (E,F) Per-base sequencing quality by recruitment source and sequencing batch, respectively. Thin curves represent individual samples and bold curves indicate group means. (G) Principal coordinates analysis of microbial community profiles, colored by recruitment source. (H) Comparison of mean species relative abundances between newly recruited participants and VinCAPR participants on logarithmic axes. Each point represents a species; the dashed diagonal indicates equal mean relative abundance. The Spearman correlation is shown, and highlighted species have *q* < 0.05.

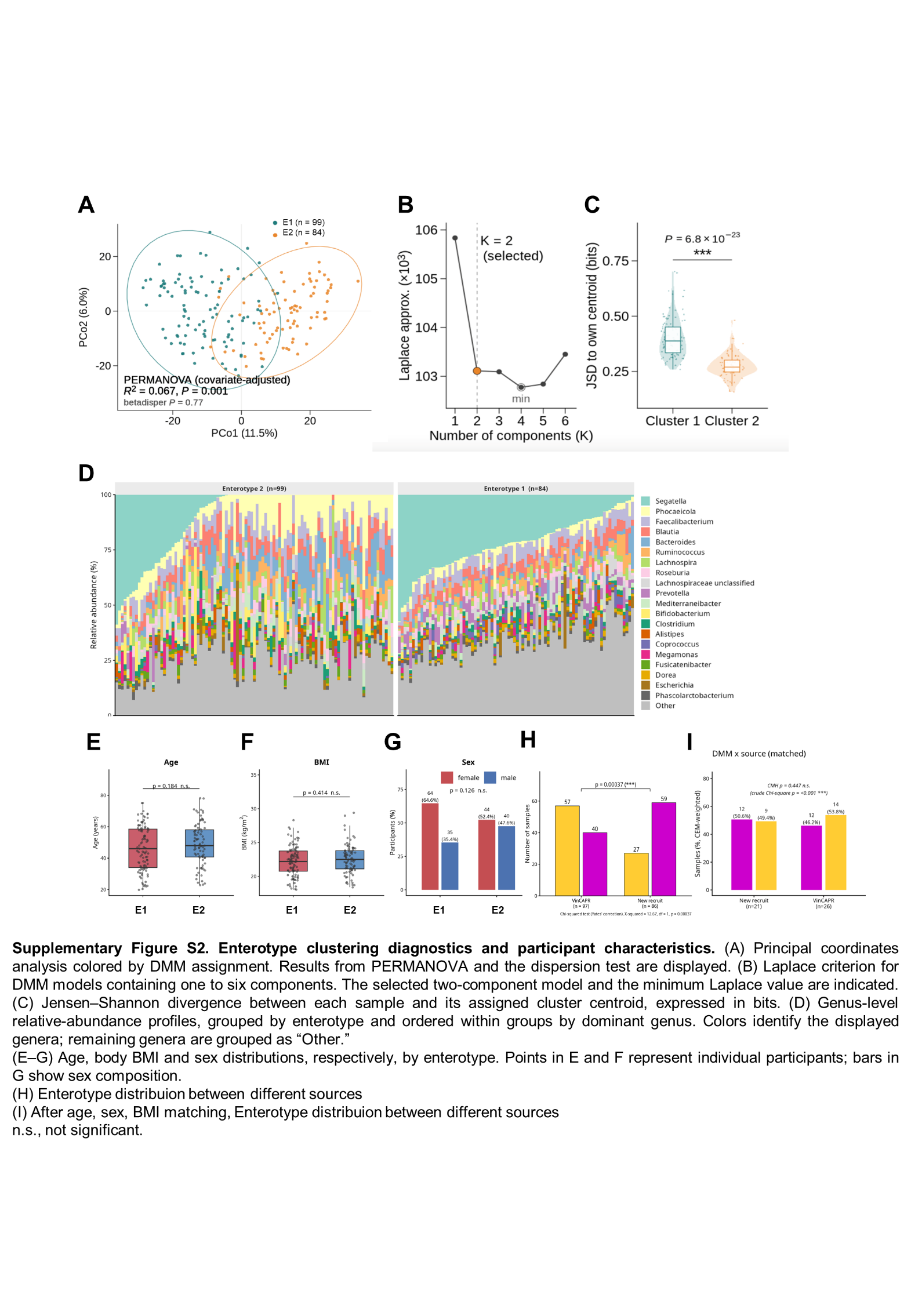

**Supplementary Figure S2. Enterotype clustering diagnostics and participant characteristics.** (A) Principal coordinates analysis colored by DMM assignment. Results from PERMANOVA and the dispersion test are displayed. (B) Laplace criterion for DMM models containing one to six components. The selected two-component model and the minimum Laplace value are indicated. (C) Jensen–Shannon divergence between each sample and its assigned cluster centroid, expressed in bits. (D) Genus-level relative-abundance profiles, grouped by enterotype and ordered within groups by dominant genus. Colors identify the displayed genera; remaining genera are grouped as “Other.” (E–G) Age, body BMI and sex distributions, respectively, by enterotype. Points in E and F represent individual participants; bars in G show sex composition. (H) Enterotype counts by recruitment source (I) Enterotype distributions by recruitment source after coarsened exact matching (CEM) on age, sex, and BMI. Bar heights represent CEM-weighted percentages. n.s., not significant; ***, *P* < 0.001

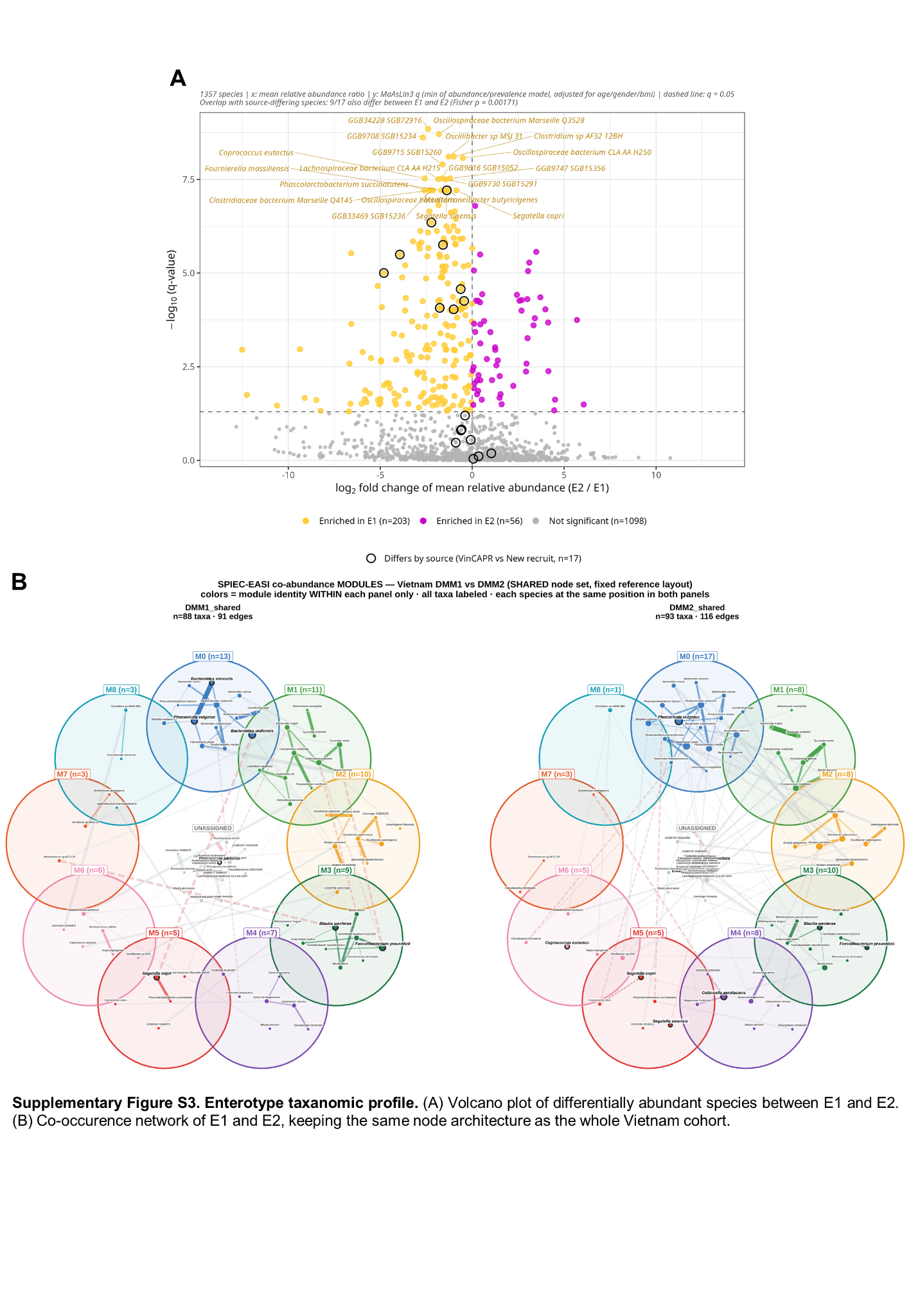

**Supplementary Figure S3. Species-level taxonomic differences and co-abundance networks by enterotype.** (A) Volcano plot of associations between species and enterotype. Yellow and magenta indicate species enriched in E1 (*n* = 203) and E2 (*n* = 56), respectively; gray indicates nonsignificant species (*n* = 1,098). Dashed lines indicate equal mean relative abundance and *q* = 0.05. Black outlines identify the 17 species that differed by recruitment source, nine of which also differed between enterotypes (Fisher’s exact test, *P* = 0.00171). (B) Species co-abundance networks inferred using SPIEC-EASI for the two enterotypes. Nodes represent species, and edges represent inferred associations. Corresponding species occupy the same positions in both networks to facilitate comparison.

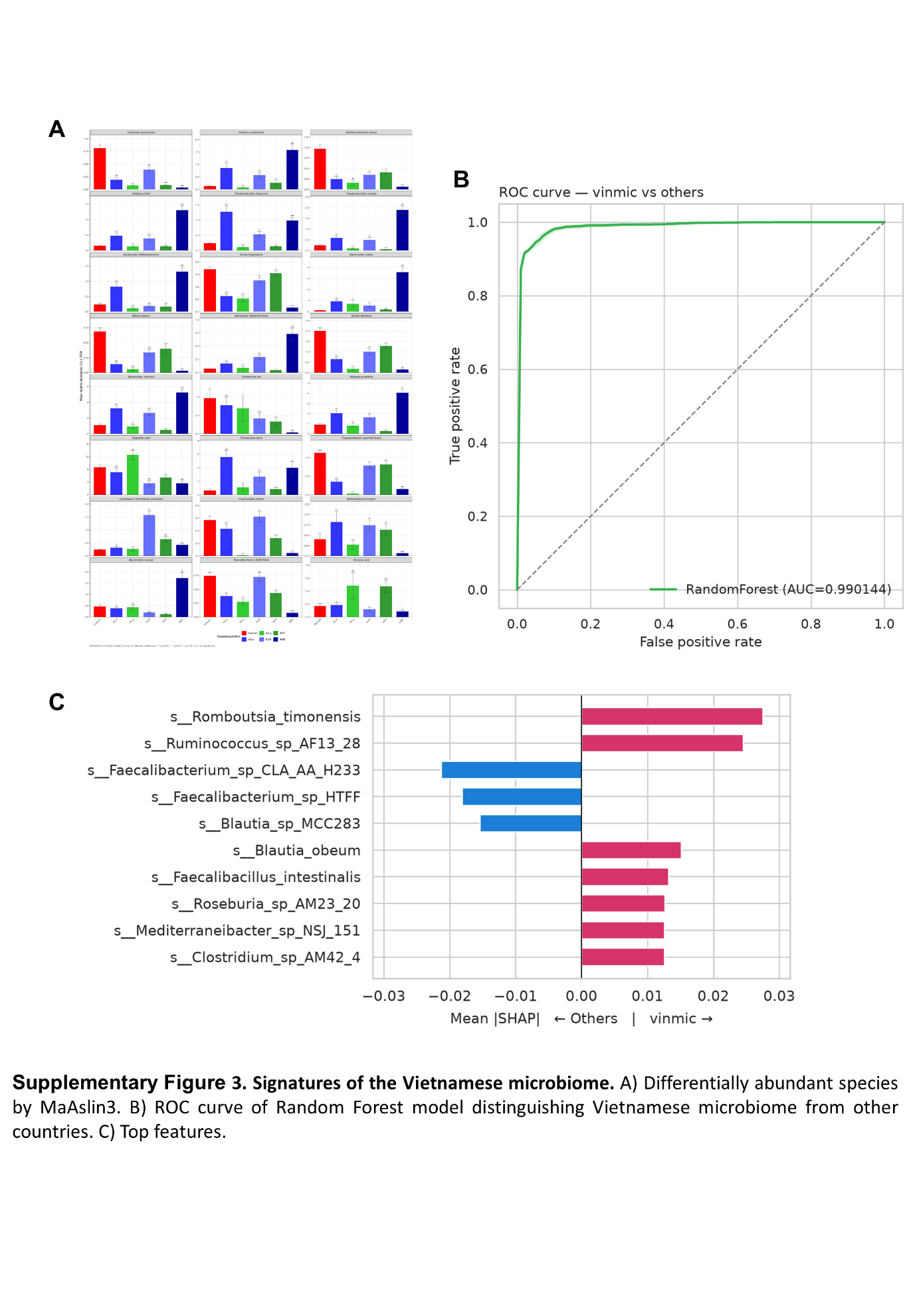

**Supplementary Figure S4. Taxonomic features distinguishing the Vietnamese cohort from public comparator populations.** (A) Relative abundances of 24 selected species across Vietnam and public comparator groups. MaAsLin3 contrasts used Vietnam as the reference and adjusted for age, sex and sequencing depth. Significance annotations indicate Benjamini–Hochberg-adjusted *q* < 0.05 (**), <0.01 (**) and <0.001 (****). (B) Receiver operating characteristic curve for a random forest classifier distinguishing Vietnamese samples from public comparator samples using core taxa (prevalence ≥20% and mean relative abundance >0.1%) (C) SHAP-based feature contributions for ten selected species, oriented toward the Vietnamese or comparator class as indicated.

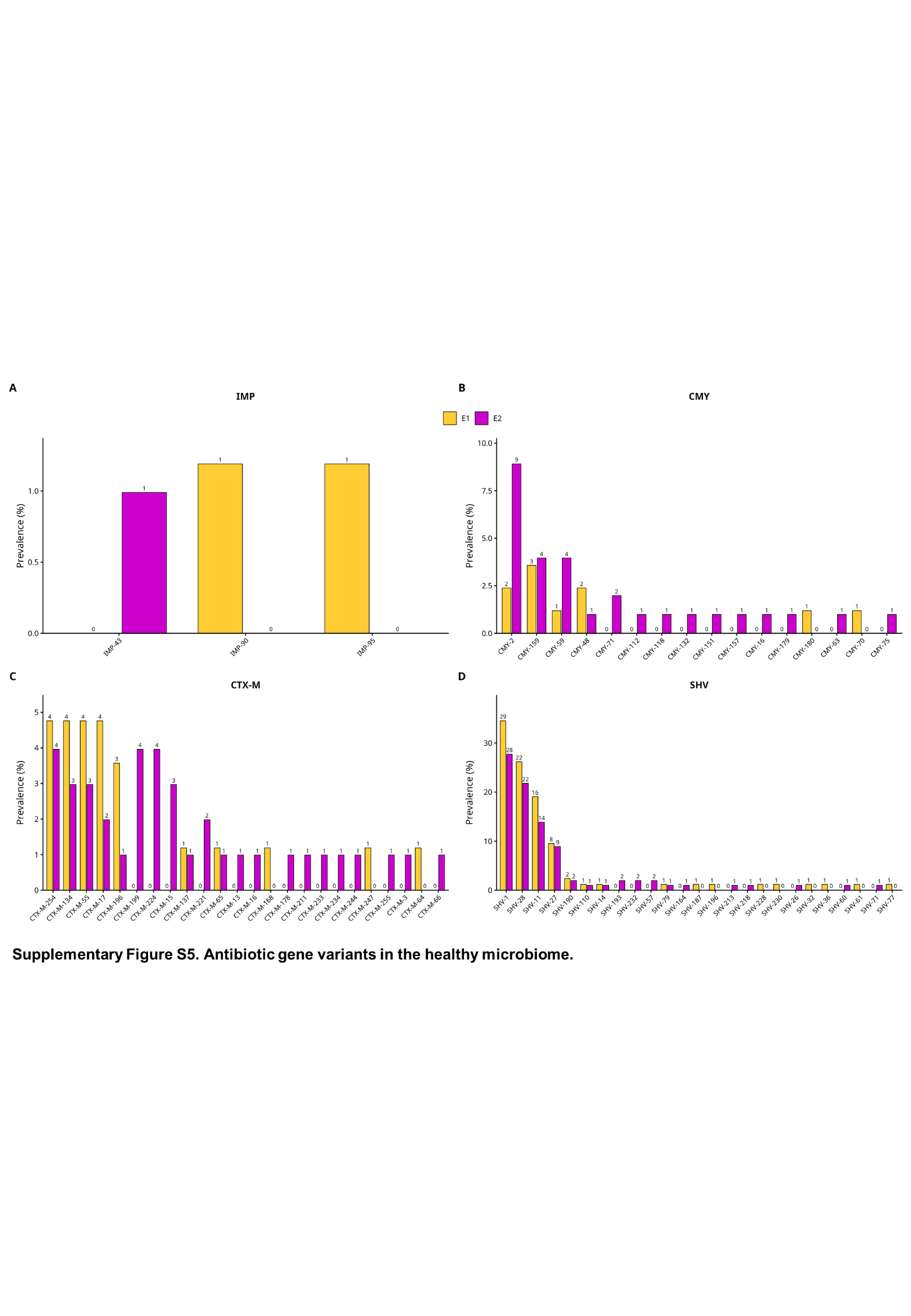

**Supplementary Figure S5. Prevalence of β-lactamase gene variants by enterotype.** Prevalence of annotated variants within the (A) IMP, (B) CMY, (C) CTX-M, and (D) SHV families in Enterotype 1 (E1; *n* = 84) and Enterotype 2 (E2; *n* = 99). Bar heights indicate the percentage of samples in each enterotype with the corresponding variant; numbers above bars indicate positive-sample counts. Yellow denotes E1 and magenta denotes E2. Comparisons are descriptive; no statistical significance is indicated.
